# Functionally annotated electrophysiological neuromarkers of healthy ageing and memory function

**DOI:** 10.1101/2023.08.26.554888

**Authors:** Tibor Auer, Robin Goldthorpe, Robert Peach, Henry Hebron, Ines R. Violante

**Affiliations:** School of Psychology, University of Surrey, Guildford, United Kingdom; Imperial College London, London, United Kingdom

**Author notes:** Corresponding author: Tibor Auer.

**Keywords:** cognitive ageing, memory, electroencephalography, connectivity, functionally annotated brain networks, pipeline

## Abstract

The unprecedented increase in life expectancy presents a unique opportunity and the necessity to explore both healthy and pathological aspects of ageing. Electroencephalography (EEG) has been widely used to identify neuromarkers of cognitive ageing due to its affordability and richness in information. However, despite the growing volume of data and methodological advancements, the abundance of contradictory and non-reproducible findings has hindered clinical translation. To address these challenges, our study introduces a comprehensive workflow expanding on previous EEG studies and investigates various static and dynamic power and connectivity estimates as potential neuromarkers of cognitive ageing in a large dataset. We also assess the robustness of our findings by testing their susceptibility to band specification. Finally, we characterise our findings using functionally annotated brain networks to improve their interpretability and multi-modal integration.

Our analysis demonstrates the effect of methodological choices on findings and that dynamic rather than static neuromarkers are not only more sensitive but also more robust. Consequently, they emerge as strong candidates for cognitive ageing neuromarkers. Moreover, we were able to replicate the most established EEG findings in cognitive ageing, such as alpha oscillation slowing, increased beta power, reduced reactivity across multiple bands, and decreased delta connectivity. Additionally, when considering individual variations in alpha band, we clarified that alpha power is characteristic of memory performance rather than ageing, highlighting its potential as a neuromarker for cognitive ageing. Finally, our approach using functionally annotated source reconstruction allowed us to provide insights into domain-specific electrophysiological mechanisms underlying memory performance and ageing.

## 1. Introduction

A major accomplishment of the 20^th^ century was the remarkable gain in global life expectancy, particularly in industrialised countries, which saw a rise of approximately 30 years (Christensen et al., 2009). However, age is also the strongest risk factor for many chronic diseases, including cancer, cardiovascular and neurodegenerative diseases (Niccoli and Partridge, 2012). Thus, understanding normal brain ageing and developing interventions that maintain cognitive function is paramount to support a better quality of life throughout one’s lifespan. Among the earliest and extensively studied cognitive changes associated with ageing is the alteration of memory (Reid and MacLullich, 2006), which also occurs in individuals ageing typically without any evidence of dementia (James et al., 2008). Accordingly, as many as half of healthy older adults’ report worrying about their everyday memory (Jonker et al., 2000).

Planning successful intervention regimes requires understanding both non-pathological and pathological changes associated with ageing and cognitive decline, alongside the identification of robust and reliable biomarkers (Belleville and Bherer, 2012; Gallen and D’Esposito, 2019; Simpraga et al., 2017). With the rise of open science and data sharing initiatives, there is now increased access to large datasets rich in phenotypic information (Collerton et al., 2007; Taylor et al., 2017), triggering several biomarker projects (Cole and Franke, 2017; Dickerson and Wolk, 2012; Engemann et al., 2020; Martin-Ruiz et al., 2011). Among the potential biomarkers, functional brain imaging metrics have increasingly emerged as promising options to monitor and identify reliable markers of brain development, ageing and disease (Foo et al., 2021; Grady, 2012; Jack et al., 2017; Nyberg et al., 2010; Tooley et al., 2022). Functional MRI has been widely used in neuroimaging studies and provides valuable information. However, it has inherent limitations, particularly as an indirect measure of neural activity, with poorer temporal resolution that hinders the characterisation of the fast dynamic properties of neural processes. In contrast, electrophysiological techniques, such as magnetoencephalography (MEG) and electroencephalography (EEG) are better suited to investigate temporal and spectral properties of neural activity across the lifespan and in the context of diseases. Recent studies have highlighted their importance in understanding the neural changes underlying Alzheimer’s Disease (AD) (Maestú et al., 2019) and their potential in neuromarker research (Engemann et al., 2020). Furthermore, EEG has the added advantage of being portable, easy to implement and an affordable imaging technique.

The high temporal resolution of M/EEG allows for the comprehensive characterisation of neural oscillations, providing insights on both normal and pathological brain function (Schnitzler and Gross, 2005). These oscillations are conventionally categorised into five classical frequency bands: delta (1-4 Hz), theta (4-7 Hz), alpha (8-13 Hz), beta (14-30 Hz), and gamma (>31 Hz). Alpha oscillations, in particular, constitute a robust electrophysiological characteristic of the awake human brain (Nunez and Srinivasan, 2009). Reduction in alpha power and alpha reactivity, defined as the reduction of alpha power upon opening the eyes, characterises not only non-pathological ageing but also pathological cognitive decline (Babiloni et al., 2022, 2006). These findings have been substantiated in a recent meta-analysis (Lejko et al., 2020) and have been associated with performance in memory, language, and executive functions (Van Der Hiele et al., 2008). Although some studies have reported evidence of age-related changes in other frequency bands, these findings are less conclusive (Trammell et al., 2017). A common limitation in previous studies is the adoption of fixed boundaries between frequency bands, which can bias findings in populations with different characteristics, such as age (Cohen, 2021). Indeed, recent studies have highlighted the importance of considering individual alpha frequency when comparing power and brain connectivity differences between younger and older adults. They demonstrate that age-related decreases in alpha frequency can bias findings in power and connectivity metrics against older adults (Jabès et al., 2021; Scally et al., 2018).

Functional connectivity measures, which commonly assess temporal correlations between time series data from two or more independent M/EEG channels or sources, have become increasingly recognised as key metrics for understanding brain regions’ communication and activity coordination, as well as identifying potential neuromarkers of ageing and disease (Engels et al., 2015; Javaid et al., 2022; Schoonhoven et al., 2022). Consequently, numerous studies have reported age-related changes in connectivity across various frequency bands, including increased connectivity in the beta band and decreased connectivity in other bands (Moezzi et al., 2019), increased connectivity in theta and beta bands, and age- and cognition-related decreases in alpha band (Chow et al., 2022). Additionally, cross-frequency coupling has shown substantial predictive power in pathological cognitive decline (Musaeus et al., 2020). However, meta-analyses revealed considerable methodological variation in functional connectivity studies, which causes significant level of inconsistency, and even contradictory findings (Lejko et al., 2020; Mahjoory et al., 2017). The vast methodological variations emphasise the importance of carefully constructed, reproducible, and shareable processing pipelines.

Furthermore, it is important to consider how we represent and summarise EEG findings to increase interpretability and facilitate comparison across different imaging modalities. In the fMRI literature, the widespread use of brain network parcellations based on structural and functional properties has contributed to the interpretability of activation patterns (Dewiputri et al., 2021). This is especially important in neuromarker research, as it strongly influences the mechanistic understanding and translational potential of the neuromarkers. While studies have demonstrated significant similarity between fMRI and M/EEG resting state networks (Brookes et al., 2011; Custo et al., 2017; Hillebrand et al., 2012), a network-based approach for reporting and interpreting the functional relevance of electrophysiological findings is scarcely used (but see (Choi et al., 2021; Zhang et al., 2021)).

In this study, we leveraged the recently published Leipzig Study for Mind-Body-Emotion Interactions (LEMON) dataset (Babayan et al., 2019), which includes, among other metrics, resting-state EEG, and memory performance assessment in over 200 young and older adult participants. Our aim was to investigate electrophysiological neuromarkers of healthy ageing and their association with memory function. Resting-state activity provides valuable insights into the intrinsic characteristics of the brain’s neural architecture (Fox and Raichle, 2007), which can reflect individual differences in cognitive function (Zou et al., 2013) and discriminate pathological conditions (Zhang et al., 2021). Resting-state data is particularly valuable in neuromarker research due to its ease of implementation, not requiring active participant engagement. This makes it more accessible and less burdensome in clinical settings, allowing for utilisation in longitudinal studies and identifying unique neurobiological signatures. Previous studies using the LEMON dataset have already begun demonstrating its utility in revealing age-related changes in brain function. For example, studies have already shown a decrease in signal variability (SD) and power in the lower frequencies (1-12 Hz) in sources related to the default mode network (DMN), as well as an age-related increase in signal variability and power in the higher frequencies (15-25 Hz and 12-30 Hz, respectively) in sources in the central frontal and temporal regions (Kumral et al., 2020; Zhong et al., 2020).

Here, we expand on previous studies conducted on the LEMON dataset and other datasets by investigating several electrophysiological neuromarkers and their association with healthy ageing and memory performance within the same participants. We included both resting state conditions, i.e., eyes open (EO) and eyes closed (EC), using two different approaches. (a) By averaging their activity (mEOEC), we increased statistical power to identify features consistent across brain states, likely corresponding to individual traits. (b) Calculating their ratio (EC/EO) as a marker of reactivity, however, can offer insights into changes in brain dynamics. This latter measure has been used in studies exploring cognitive decline (Barry and De Blasio, 2017) and predicting cognitive performance (Van Der Hiele et al., 2008). We investigated power and functional connectivity, both within- and across frequency bands, for both approaches. Given the growing recognition of individualised frequency bands, we considered how individual variations affected age- and memory-related findings on activity and connectivity metrics. All analyses were performed at the source level, thus obtaining a more fine-grained topographic distribution of the features, which is important in predicting cognitive ageing (Engemann et al., 2020). Finally, we interpret the results on the level of neuropsychologically meaningful networks to allow a more integrative view and a more direct link between brain and behaviour. To support open and reproducible science, we implemented our analyses using a configurable and scalable neuroimaging pipeline (Cusack et al., 2015).

## 2. Methods

### 2.1. Participants

This study used data from the Leipzig Study for Mind-Body-Emotion Interactions (LEMON) project (Babayan et al., 2019), including 227 participants (82 female) divided into nine pre-defined age groups with age ranges centred at 22.5, 27.5, 32.5, 37.5, 57.5, 62.5, 67.5, 72.5, and 77.5. Participants were further labelled as “young” (20-40 years old) and “older” (55-80 years old). Exclusion criteria included ongoing substance misuse, neurological disorders, malignant disease, cardiovascular disease, psychiatric illness requiring inpatient treatment, pregnancy, claustrophobia, metallic body implants including tattoos, tinnitus, hypertension, recent involvement in research or advanced psychology degrees, and certain medications including those acting on the central nervous system, chemotherapy, and psychopharmacological medicines. Data collection was in accordance with the Declaration of Helsinki and was approved by the medical faculty ethics committee at the University of Leipzig (reference 154/13-ff).

### 2.2. Behavioural data

Participants completed a cognitive test battery and the resting state (rs)-EEG on separate sequential daily sessions. Out of test battery, we selected the two tests that directly measured memory function, i.e., an adapted version of the California Verbal Learning Test (CVLT-II, (Niemann et al., 2008)) and the 2-Back task from the Test of Attentional Performance (TAP; (Ziemmermann and Fimm, 2021, n.d.)). The CVLT-II was done first, followed by the TAP as a 20-minute filler task, and the CVLT-II delayed recall condition after. TAP was employed as a filler task as it is unlikely to interfere with verbal learning (Jones et al., 2014).

The CVLT-II was used to assess participants’ verbal learning and episodic memory. Participants were acoustically presented with 16 words and instructed to remember them. They were then presented with this same list and asked to immediately recall its contents five consecutive times. This was followed by several different recall conditions. In the interference condition, participants learned a new list once and immediately recalled the original list. In the cued condition, participants were presented with four categories and asked to recall which words from the original list correspond with which category. In the delayed condition, participants were asked to recall the original list after a delay of 20 minutes, both with and without category cues. In the recognition condition, participants were presented with a new list of words and asked to determine which of these were on the original list. Cued conditions were used to assess associative memory functioning, whilst performance across the other conditions assessed episodic memory.

The 2-Back task from the TAP was used to assess working memory. This task, presented visually to participants on a computer screen, serially showed a list of numbers (1-9) for 5 minutes. Participants were required to press a button if the number currently on the screen matched the one they saw two numbers prior. Accuracy, omissions, and reaction times were recorded for each participant to assess their performance.

Composite measures of associative (AM), episodic (EM), and working memory (WM) were created by extracting scores from the CVLT-II and TAP. These composite scores were favoured over individual test scores to allow for a more comprehensive memory assessment of each domain. A composite episodic memory score was created by factor analysing CVLT-II recall on first and fifth trials, the sum of correct recalls from the first to fifth trial, recall after a short delay, delayed recall by 20 minutes, and recognition of learned words. All measures correlated above 0.4, Kaiser-Meyer-Olkin’s measurement of sampling adequacy considerably exceeded 0.5 at 0.81, and Bartlett’s test of Sphericity was significant (*Х^2^*(15) = 1273.51, *p* < 0.001), all of which indicated good factorizability. From this, one principal component with an Eigenvalue above 1 emerged, which accounted for 71.24% of the variance across episodic memory scores. A composite associative memory score was similarly created by factor analysing word recall after a short delay with category cues present and delayed recall by 20 minutes with category cues present. Measures were highly correlated, given the smaller number of variables, while the Kaiser-Meyer-Olkin coefficient was lower but still acceptable at 0.5 (Hadia et al., 2016), and Bartlett’s test of Sphericity was significant (*Х^2^*(1) = 425.50, *p* < 0.001). From this analysis, one principal component with an Eigenvalue > 1 emerged, which accounted for 96.16% of the variance across associative memory scores. An index score was created to assess working memory faculties measured with the accuracy on the 2-Back TAP task. This index was calculated by subtracting the number of incorrect responses from the number of correct responses and dividing the total by 15 (the number of possible correct responses).

### 2.3. EEG

#### 2.3.1. Data acquisition

Sixteen minutes of rs-EEG data were acquired for 216 participants using a BrainAmp MR plus amplifier with 62-channel active ActiCAP electrodes (Brain Products GmbH, Germany). Electrodes were placed according to the 10-10 localisation system, referenced to the FCz electrode, and the ground electrode was placed on the sternum. Skin electrode impedance was kept below 5 KΩ. During EEG data acquisition, EEG amplitude resolution was set to 0.1 μV, data was digitised at a sampling rate of 2500 Hz, and recorded with an online band-pass filter between 0.015 Hz and 1 kHz.

Rs-EEG data was collected over 16 blocks, each lasting 60 seconds. Blocks were divided into eight eyes-open (EO) and eight eyes-closed (EC) conditions interleaved. Changes between blocks were announced using Presentation software (version 16.5, Neurobehavioral Systems Inc., Berkeley, CA, USA). Participants were seated in front of a computer screen and asked to stay awake during EEG data acquisition. During the EO blocks, they were asked to fixate their gaze on a black cross presented on a white background.

#### 2.3.2. Pre-processing

We opted for using the raw data instead of the available pre-processed EEG data for two main reasons. The first was that the published pre-processed data had been band-passed with 1-45 Hz, thus removing information on high-frequencies which may hold important information regarding cognition and age-related changes in brain function (Başar, 2013; Herrmann et al., 2010, 2004). The second was that this allowed us to implement an open access processing workflow, which allows us better control over data quality, larger transparency, and flexibility in pre-processing.

Raw rs-EEG was pre-processed using Automatic Analysis (*aa*, version 5.6; https://automaticanalysis.github.io, (Cusack et al., 2015)) running on MATLAB R2020a (Mathworks, Inc, Natick, Massachusetts, USA). The workflow (**Fig. S1**) included pre-processing using EEGLab (version 2020.0; (Delorme and Makeig, 2004)) and FieldTrip (git revision 666b4e3; (Oostenveld et al., 2011)). Raw rs-EEG data was initially down-sampled from 2500 Hz to 250 Hz and high-pass filtered with 1 Hz. Line noise was removed using a band-pass filter at 50 Hz and 100 Hz with a bandwidth of 10 Hz. Artefactual channels and data segments were removed using Artifact Subspace Reconstruction (Chang et al., 2018), and data were re-referenced to a common average reference before further data processing. Further pre-processing included Independent Component Analysis using the AMICA algorithm (Delorme et al., 2012; Palmer et al., 2007) followed by automated IC selection using IClabel (Pion-Tonachini et al., 2019) and dipole fitting to inform classification of components. Finally, data were divided into EO and EC conditions and epoched with a 2-second interval within each condition (maximum eight bock x 30 = 240 epochs). Following pre-processing, 163 participants’ EEG data with at least 100 clean epochs were retained for further analysis.

#### 2.3.3. Data analysis

Data analysis was conducted using FieldTrip (git revision 666b4e3; (Oostenveld et al., 2011)) as integrated into *aa* (**Fig. S1**). Power spectral density (PSD) was calculated using fast Fourier transforms across the spectrum between 1 Hz and 120 Hz after ‘tapering’ the data with the Hanning window. Across all participants, a standard, highly-detailed Finite Element Method volume conduction model was used to solve the forward problem (Huang et al., 2016) using the SimBio toolbox as integrated into FieldTrip (Vorwerk et al., 2018). The source model was created based on the cortical sheet of each participant as constructed with FreeSurfer and downsampled to around 4000 tessels using Connectome Workbench (https://www.humanconnectome.org/software/connectome-workbench). Source activity was reconstructed by using exact low-resolution brain electromagnetic tomography (eLORETA) as implemented in FieldTrip (Pascual-Marqui, 2007). The leadfield matrix and the source filter were generated between the modelled cortical sources and the EEG channels and were used to compute the abovementioned time-frequency decomposition at the source level.

The epoched signal was computed at the source level and averaged for regions of the Desikan-Killiany-Tourville (DKT) atlas (Desikan et al., 2006). The Freesurfer parcellation annotated 62 regions according to the DKT atlas; however, 11 of them (the entorhinal, the cingulate isthmus, the medial orbitofrontal, the parahippocampal, and the pars orbitalis of the inferior frontal bilaterally, as well as the right posterior cingulate) failed to be mapped to the source model thus leaving 51 virtual channels. Band-limited power spectrums were calculated by averaging the PSD according to standard EEG frequency bands (delta: 1-3 Hz; theta: 4-7 Hz; alpha: 8-13 Hz; beta: 14-32 Hz; lower gamma: 33-80 Hz; upper gamma: 81-120 Hz) before statistical analysis. For connectivity, time-frequency decomposition was averaged according to 33 bins with increasing width (delta: 6 bins; theta: 7 bins; alpha: 6 bins; beta: 5 bins; lower gamma: 5 bins; upper gamma: 4 bins) and two measures were computed between virtual channels. These were the within-frequency connectivity, as characterised by the debiased weighted phase lag index (wPLI) (Vinck et al., 2011), and cross-frequency connectivity as characterised by the phase locking value (PLV) (Schmidt et al., 2014). In a recent study, Siebenhühner and co-workers demonstrated the reliability and biological plausibility of these measures (Siebenhühner et al., 2020), as well as their correspondence with cognitive performance.

In addition to considering standard EEG frequency bands, we estimated individual bands using a combination of extended Better OSCillation detection (eBOSC) (Kosciessa et al., 2020) and Fitting Oscillations & One Over F (FOOOF) (Donoghue et al., 2020). eBOSC uses a 6-wave wavelet transform across the spectrum to calculate PSD for individual band detection. On the other hand, FOOOF operates by parameterising a PSD model by fitting Gaussian curves to capture band-limited power spectra as peak-like deviations from the background activity. After calculating the PSD across the spectrum between 2 Hz and 80 Hz, a FOOOF model was fitted to detect up to six Gaussian curves with a peak width between 1 and 6 Hz and a minimum peak height of 0.05 a.u. or 1.5 SD, whichever is higher. The band estimates (peak frequency and bandwidth) were averaged across channels to calculate the individual bands. The procedure provides stable estimates for alpha and beta bands within the standard bands (see above). Theta band was shifted accordingly, while keeping its original bandwidth; however, delta band’s width was adjusted to correspond to all frequencies below the theta band. Gamma bands were unaffected. Individual variations in band frequencies are accounted for in all results reported in the main text, while corresponding results without band individualisation are reported in the supplemental material for comparison.

#### 2.3.4. Combined neural measures

Averaging measures for EC and EO conditions (mean (m) ECEO, calculated as 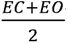) allows the investigation of neural features stable across the two states. The literature on defining EC/EO reactivity is not conclusive, and various approaches have been reported from simple difference (Bellato et al., 2020), through ratio (Fonseca et al., 2011), to normalised difference (Wan et al., 2019); usually without detailed justification. Considering a report of a linear relationship between EC and EO estimates in all frequency bands (Barry and De Blasio, 2017) also seen in our data (*not shown*), we decided to use their ratio 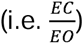 as a measure of EC/EO reactivity.

The combinations of neural measures have been performed at the last stage of the analysis before statistics. For example, source-level power estimates have been calculated for the EC and the EO conditions separately, and then these source-level estimates have been combined (average and ratio, respectively) to obtain the final features we entered in the statistical analysis.

#### 2.3.5. Statistical analysis

Behavioural data was analysed with R version 3.6.0 (2019-04-26). Since none of the behavioural data showed a normal distribution (AM: W = 0.94, p < 0.001; EM: W = 0.97, p = 0.003; WM: W = 0.81, p < 0.001), we used Mann–Whitney U tests and Kruskal–Wallis tests when comparing “young” and “older” groups and the nine pre-defined age groups, respectively. Effect sizes have been interpreted according to Funder and Ozer (Funder and Ozer, 2019).

The effect of age and the neural correlate of the associative, episodic, and working memory domains were tested on all estimates, i.e., the time-frequency decomposition and the within- and cross-frequency connectivity. The effect of age was tested by means of linear regression using the interval variable of age groups as the independent variable. The neural correlates of memory domains were tested by means of linear regression using the composite measures as the independent variable. Due to the strong linear relationship between age and memory performances (see Results), these linear regressions have been conducted for the “young” and “older” groups separately. Also, the effect of age on memory performances has been accounted for by the orthogonalisation of memory performances with respect to age within each group.

Statistical inference was calculated by means of nonparametric Monte-Carlo estimation of the significance probabilities as implemented in FieldTrip. For all estimates, statistical significance was calculated based on 1000 iterations of threshold-free cluster enhancement (TFCE), and a significance threshold of p = 0.05 was employed. TFCE- and other cluster-based statistics are robust inferential procedures offering greater freedom and sensitivity (Maris and Oostenveld, 2007). They, however, require clear rules on cluster formation, i.e., which data points can form a cluster. If we consider time-frequency decomposition, where the data is acquired from single channels, a cluster-forming rule can be defined based on the topographic distribution of the channels based on the assumption that data from channels spatially closer to each other are likely to be more similar (Frömer et al., 2018). However, defining a cluster-forming rule for channel combinations for connectivity is less straightforward since they lack a clearly defined topographic property (both channels correspond to their spatial locations).

#### 2.3.6. Graphs for cluster-forming rules for functional connectivity

To address the issue of creating cluster-forming rules for channel combinations for connectivity metrics we used graphs, i.e., in a similar fashion to how channels and their connectivity (spatial and functional) can be modelled as graphs, channel combinations can also be modelled as graphs. Our rationale and procedure is described below.

Let us begin with a standard graph where the nodes represent channels, and the edges link any pair of channels either spatially (*G*_*spat*_, **Fig. 1A**) or via hypothesised functional connectivity (*G*_*conn*_, **Fig. 1B**). Adjacencies between the edges of a graph *G* can be found using the line-graph representation *L(G)*, where each node in *L(G)* corresponds to an edge in *G* (Harary and Norman, 1960) (*L(G*_*spat*_ ) and *L(G*_*conn*_), Fig 1C-D). More explicitly, if two edges in *G* are incident on a common node, they will be connected via an edge in *L(G)*, i.e. they will be adjacent. A further restriction on the edge-adjacency in an *L(G)* can be applied, whereby two edges can only be adjacent if the non-shared nodes are also adjacent in *G*. This additional restriction on edge-adjacency is based on the rationale that extends the aforementioned observation of data similarity to connections, namely, connections between a channel to a set of channels adjacent to each other are likely to be more similar. The result of this additional restriction is the restricted line graph *L*_*r*_*(G)*, and we can have one for the spatial connectivity (*L*_*r*_*(G*_*spat*_ ), **Fig 1E**) and another for the hypothesised functional connectivity (*L*_*r*_*(G*_*conn*_), **Fig. 1F**).

**Figure 1.**
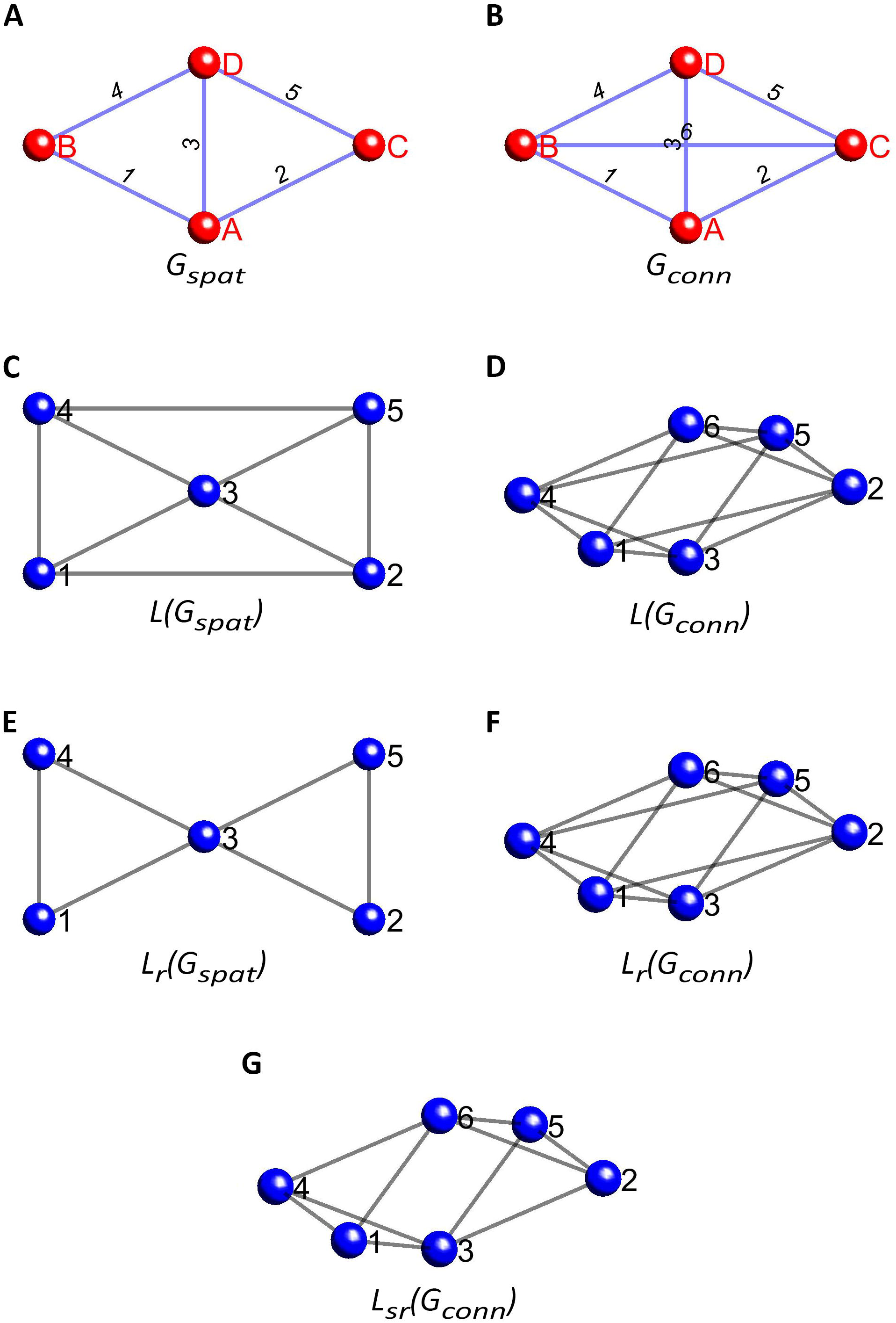
Clustering of connectivity based on graph theory. **(A)** Hypothetical layout of signal visualised as a *G*_*spat*_. The red nodes marked with upper-cased letters correspond to the signal locations, while the blue edges marked with numbers correspond to the spatial relationship of the signal locations. **(B)** Connectivity to be tested for the signals in *G*_*spat*_ visualised as a *G*_*conn*_. The red nodes marked with upper-cased letters correspond to the same signal locations as in *G*_*spat*_, while the blue edges marked with numbers correspond to the functional connectivity to be tested. **(C)** and **(D)** represents the line-graphs of *G*_*spat*_ and *G*_*conn*_, respectively. The blue nodes correspond to the edges of original graphs, while the grey lines correspond to the egde-adjacencies (i.e., neighbourhoodness of edges). **(E)** and **(F)** represents the line-graphs of *G*_*spat*_ and *G*_*conn*_, respectively, with adjacency restricted to edges with adjacent non-shared nodes. As you can see, edges 4 and 5 of *G*_*spat*_ are not adjacent anymore because their non-shared nodes (B and C) are also not adjacent. It does not change the line-graph of *G*_*conn*_ because all connectivity between all signals is of our interest and, therefore, all nodes are connected to all nodes. **(G)** represents the final solution, where the edge-adjacency for *G*_*conn*_ is restricted based on the node-adjacency in *G*_*spat*_. As a result, edges 4 and 5 of *G*_*conn*_ are not adjacent anymore.

This additional restriction of edge-adjacency can be generalised so that the restricted edge-adjacency of a hypothesised functional connectivity graph *L*_*r*_*(G*_*conn*_) is based on the node-adjacency in a spatial graph *G*_*spat*_. More explicitly, two edges can only be adjacent in *L*_*r*_*(G*_*conn*_ ) if the non-shared nodes are also adjacent in *G*_*spat*_. This concept is demonstrated in **Fig. 1G**, wherein the line-graph of the fully connected hypothesised functional connectivity *L(G*_*conn*_) is restricted by the spatial graph *G*_*spat*_ to return a spatially restricted line graph *L*_*sr*_*(G*_*conn*_). Looking closely, one can see that nodes 1 and 2 of *L*_*sr*_ *(G*_*conn*_) (which are edges in the original hypothesised functional connectivity graph *G*_*conn*_), are not connected because even though they share node A in *G*_*conn*_, the non-shared nodes (B and C) are not adjacent in the spatial graph *G*_*spat*_. For inferencing on connectivity, the cluster-forming rule is defined based on the spatially restricted line graph *L*_*sr*_ *(G*_*conn*_ ).

## 3. Results

### 3.1. Demographics and behavioural data

The final sample included 163 participants, 112 (33 female) in the “young” group and 51 (24 female) from the “older” group. The sample distribution across the nine age groups can be observed in **Table S1** and **Fig. 2A**.

**Figure 2.**
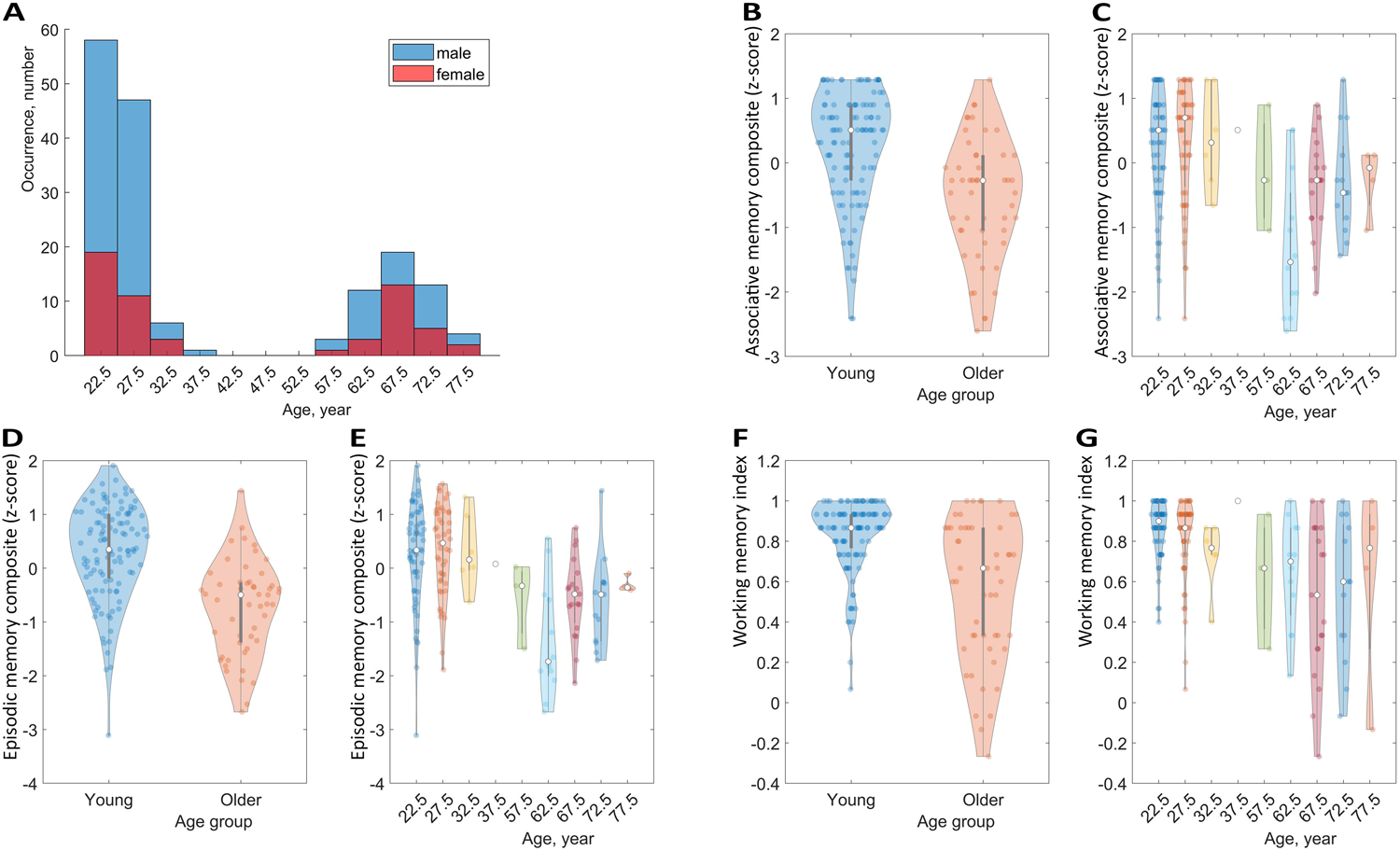
Demographics and memory performance of the final sample. **(A)** Distribution of age and gender. **(B)**, **(D)** and **(F)** show the distribution of performance in associative (B), episodic (D) and working (F) memory for the participants labelled as “young” and “older”. **(C)**, **(E)** and **(G)** show the distribution of performance in the same memory domains for the participants in the original age groups. In plots **(B-G)**, the dots represent individual performances.

The composite memory scores are displayed in **Fig. 2B-G** and **Table S2** for both the “young” and “older” groups, and for the nine age groups. Wilcoxon rank sum tests were performed to compare the composite associative (AM), episodic (EM), and working (WM) memory scores between the “young” and the “older” group. The composite memory scores were significantly larger in younger than older adults’ groups in every domain, with a very large effect size, AM (**Fig. 2B**): W=4233.5, p<0.001, r = 0.48; EM (**Fig. 2D**): W=4591, p<0.001, r = 0.61; WM (**Fig. 2F**): W=4235.5, p<0.001, r = 0.48. Considering the nine pre-defined age groups the Kruskal-Wallis rank sum test also resulted in significant differences with small to medium effects, AM (**Fig. 2C**): *Х*^2^(8)=28.61, p<0.001, r = 0.18; EM (**Fig. 2E**): *Х*^2^(8)=40.99, p<0.001, r = 0.25; WM (**Fig. 2G**): *Х*^2^(8)=31.18, p<0.001, r = 0.19.

### 3.2. Age-related changes in mEOEC power and EO/EC reactivity and relation to memory performance

#### 3.2.1. Age-related changes in frequency and the effects of band individualisation on power

We first investigated whether individual peak frequencies within alpha and beta frequency ranges varied with age. The topographic distribution of the effect of ageing revealed global changes in peak frequency for both alpha and beta frequencies, with somewhat weaker effect in the frontal regions for alpha and the posterior midline regions for both alpha and beta (**Fig. 3**, topoplots). This effect was statistically significant only for alpha in the EC condition in the frontal midline, occipital, and right temporal regions. The analysis of peak frequencies revealed an age-related downward shift in individual alpha and beta across sensors (**Fig. 3**, regression plots, data was averaged across all sensors for each individualised frequency band). This shift was larger in the EC conditions for frequencies in the alpha range (EC: 0.008 Hz/year, EO: 0.002 Hz/year) and in the EO condition for frequencies in the beta range (EC: 0.01 Hz/year, EO: 0.019 Hz/year), reaching significance for alpha EC (t(161)=-2.294, p=0.023) and near-significance for beta EO (t(161)=-1.935, p=0.055). This indicates that the effects of ageing on peak frequency changes are more robustly detected in the alpha band for the EC condition. This might in part be explained by the relatively larger signal-to-noise ratio in alpha during EC. Importantly, there is a large between-subject variance, which is even larger than the effect of ageing, which explains only 2-3% of the variance in the data. These results highlight the importance of condition-specific individualisation beyond simply scaling with age.

**Figure 3.**
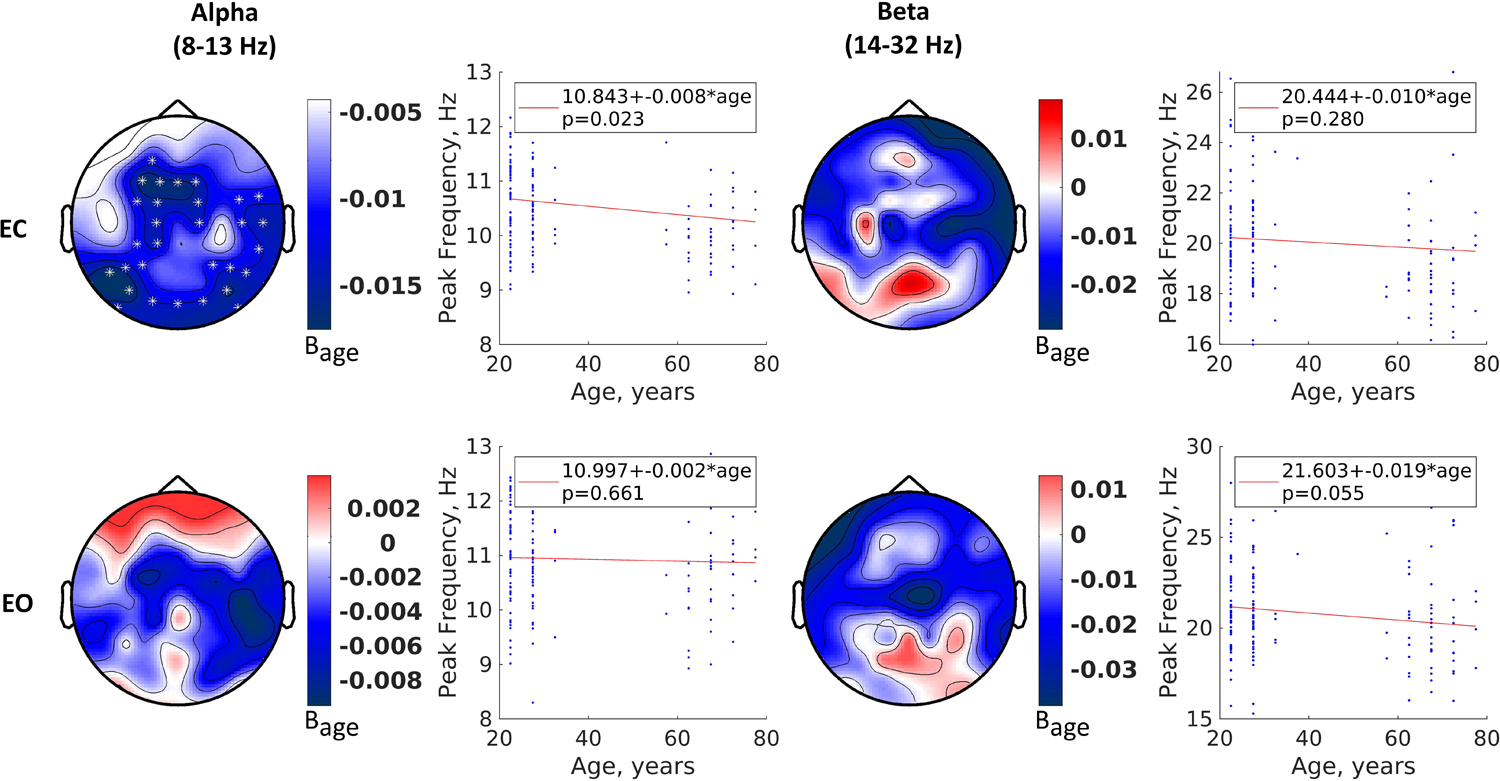
Age-related changes in alpha and beta peak frequency. The topoplots demonstrate the localised effect of age on the peak frequencies for alpha and beta and for eyes-closed (EC) and eyes-open (EO) conditions. The values correspond to the slope (B) of the regression analysis. The white stars correspond to significant effect. The line and scatter plots demonstrate the effect of age on the same peak frequencies averaged over all channels. The red lines correspond to the regression line, while the blue dots correspond to the individual peak frequencies.

Next, we investigated the effect of individualised frequency ranges on power estimates by comparing the relative differences observed between the individually determined and canonical bands. Individualised alpha and beta bands were centred at the individual alpha and beta peak frequencies ± the estimated half-bandwidth, delta and theta bands were shifted to ensure there was no overlap between the bands while keeping their canonical bandwidth. Compared with standard bands, band individualisation resulted in around 10% global reduction in mEOEC power with more emphasis on the frontal and central areas in the delta and theta power and a 10% localised increase in the temporo-occipital areas in the theta power (**Fig. 4**, top left half). On the other hand, band individualisation strongly increased the alpha and beta power, resulting in up to 18% and 110% increase, respectively, with a maximum effect in the frontal areas (alpha) an the temporo-parieto-occipital junction (beta) (**Fig. 4**, top right half). The band individualisation also affected the reactivity in power, although with a smaller effect. There was an up to 2% global increase in delta reactivity with more emphasis on the central regions, while there was an up to 7% increase in theta reactivity in the occipital regions (**Fig. 4**, bottom left half). The alpha and beta reactivity increases were larger, up to 10% and 30%, respectively, and primarily included the frontal and posterior regions (**Fig. 4**, bottom right half). Overall, these results support the importance of considering individually defined frequency bands when exploring age-related changes in brain function.

**Figure 4.**
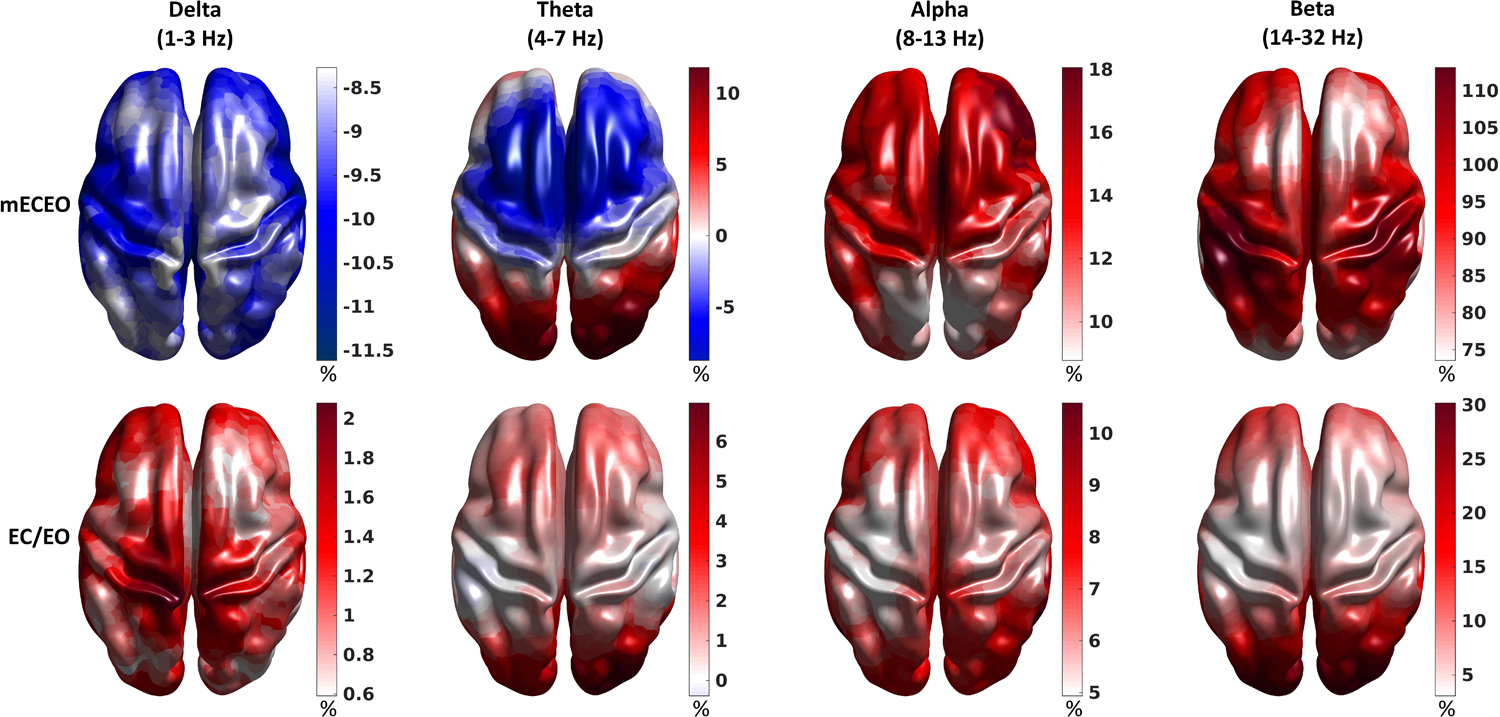
The effect of band individualisation on the combined neural measures. The surface plots demonstrate the localised effect of band individualisation on the combined neural measures of mean power (mECEO) and reactivity (EC/EO). The values correspond to how much the measures change (in % of the measures estimated in the canonical bands) after individualisation.

#### 3.2.2. Age-related changes in mean power and reactivity

Our results for mean power and power reactivity are displayed in **Fig. 5** for individualised bands and in **Fig. S2** for canonical frequency bands; **Table 1** summarises the age-related changes per frequency band (individualised and canonical) for each brain network.

**Figure 5.**
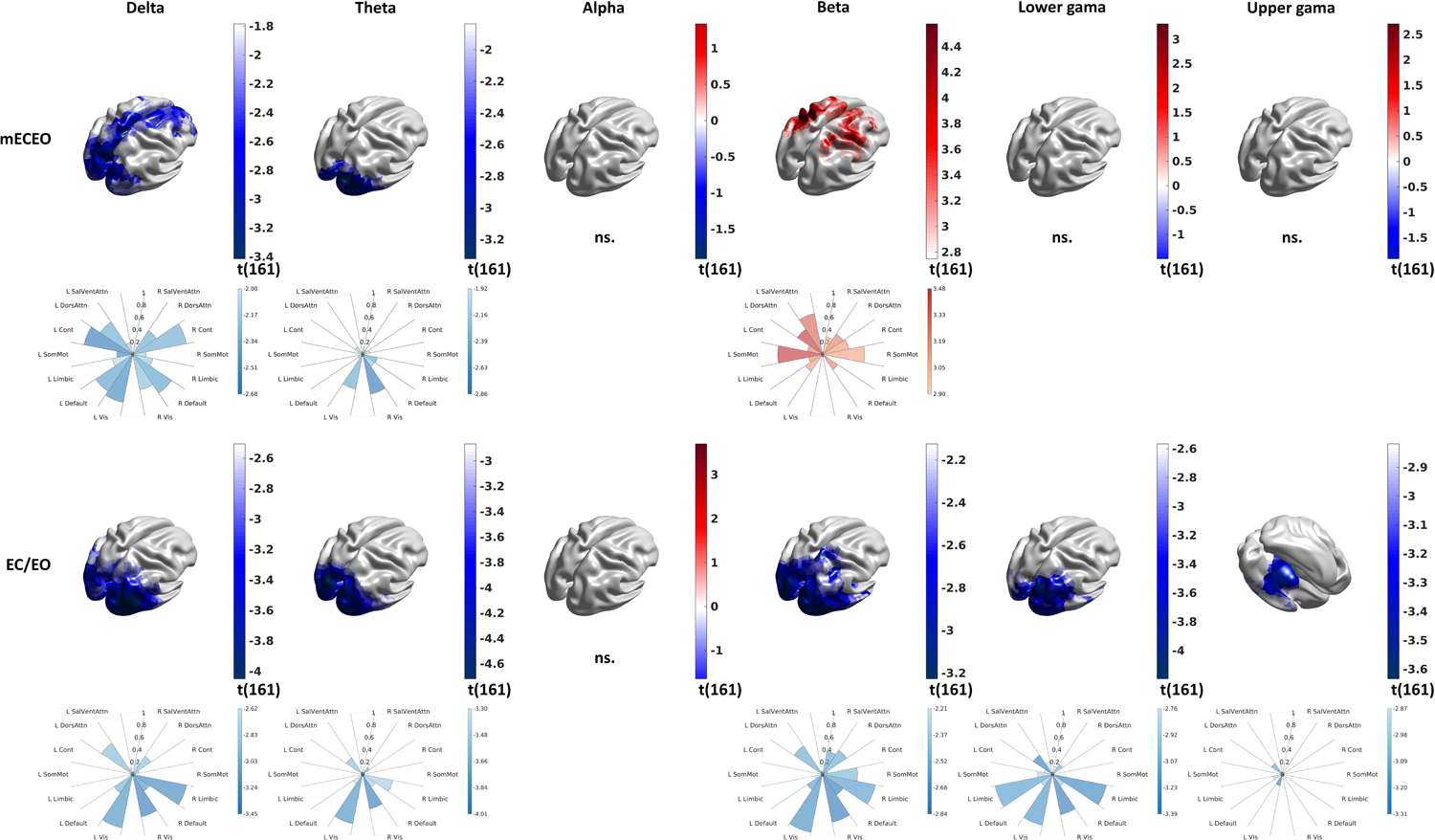
Age-related changes in the combined neural measures. The surface plots demonstrate the significant localised effect of ageing, measured as t-statistics of the regression, on the combined neural measures of mean power (mECEO) and reactivity (EC/EO) in the individualised bands. “ns.” denotes cased with no significant effect. The polar plots visualise the detected effects averaged in the seven functionally annotated networks in both hemispheres. The colour corresponds to the effect size as measured with the t-statistics of the regression, while the size of the wedges corresponds to the proportion of the functional network involved.

**Table 1.**
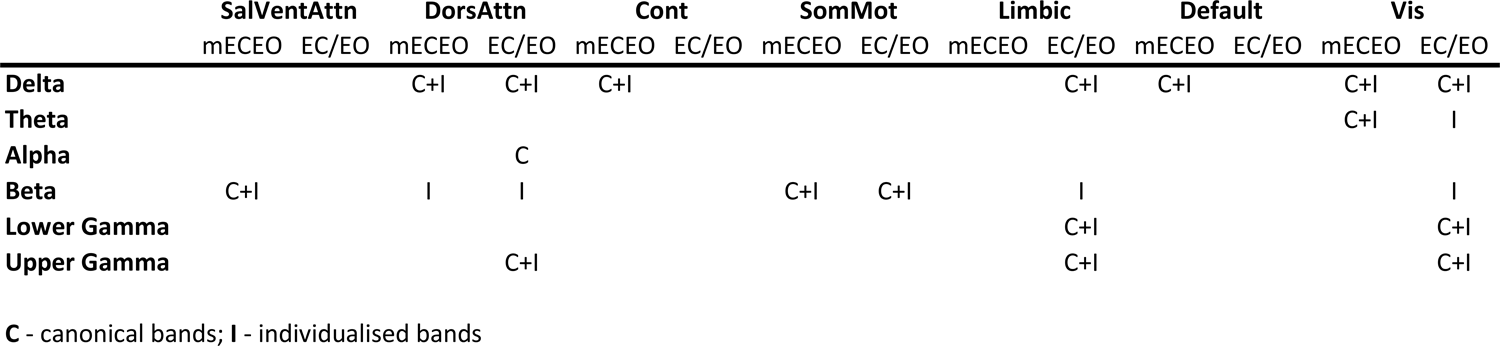
The effect of age and band individualisation on the combined neural measures. The table summarises in which functionally annotated networks we detected age-related changes in mean power (mECEO) and reactivity (EC/EO) by using canonical (C) and individualised (I) bands.

We observed an age-related reduction in mEOEC delta power in the occipital and midfrontal regions corresponding to the Visual, Default, Control, and Dorsal Attention networks. Delta power reactivity showed more focal changes in the Visual, Dorsal Attention, and Limbic networks. Here, band individualisation made only minor difference (**Fig. 5** vs **Fig. S2**). We found age-related reduction of mean theta power and reactivity in the occipital regions corresponding to the Visual network and no changes in the midline regions. Here, band individualisation strongly increased the sensitivity to detect the ageing effect in theta reactivity. We saw no effect in alpha power or reactivity. Here, band individualisation made only minor difference by pushing a very focal effect of ageing below the significance threshold in alpha reactivity in the right prefrontal region. Aging affected beta mEOEC power and reactivity differently, increasing mEOEC beta power in the Somatomotor and Dorsal and Ventral Attention networks while reducing beta reactivity in the Visual, Limbic, Somatomotor, and Dorsal Attention networks. Band individualisation strongly increased the sensitivity to detect ageing effects in beta power. We saw an age-related reduction in reactivity in the lower gamma band in the Visual and Limbic networks. Finally, we observed some age-related reductions in reactivity in the upper gamma band in the Visual, Dorsal Attention, and Limbic networks.

#### 3.2.3. Mean power and reactivity correlates of memory

Analyses of the relationship between memory performance and the mean power estimates only showed significant correlations with performance in working memory for both young and older adults (**Fig. 6**). mEOEC power in individualised bands was positively correlated with working memory performance in the alpha band for both young and older adults. The topography showed overlapping left-lateralised correlation in both groups in regions belonging to the Dorsal Attention network, occupying a larger area in the young group (**Fig. 6A** and **Table 2**). Furthermore, younger adults also showed significantly positive correlations with a similar topography for lower and upper gamma activity. Overall, this supports previous findings of the importance of the Dorsal and Ventral Attention networks and alpha and gamma activity in working memory (Jokisch and Jensen, 2007; Majerus et al., 2018).

**Figure 6.**
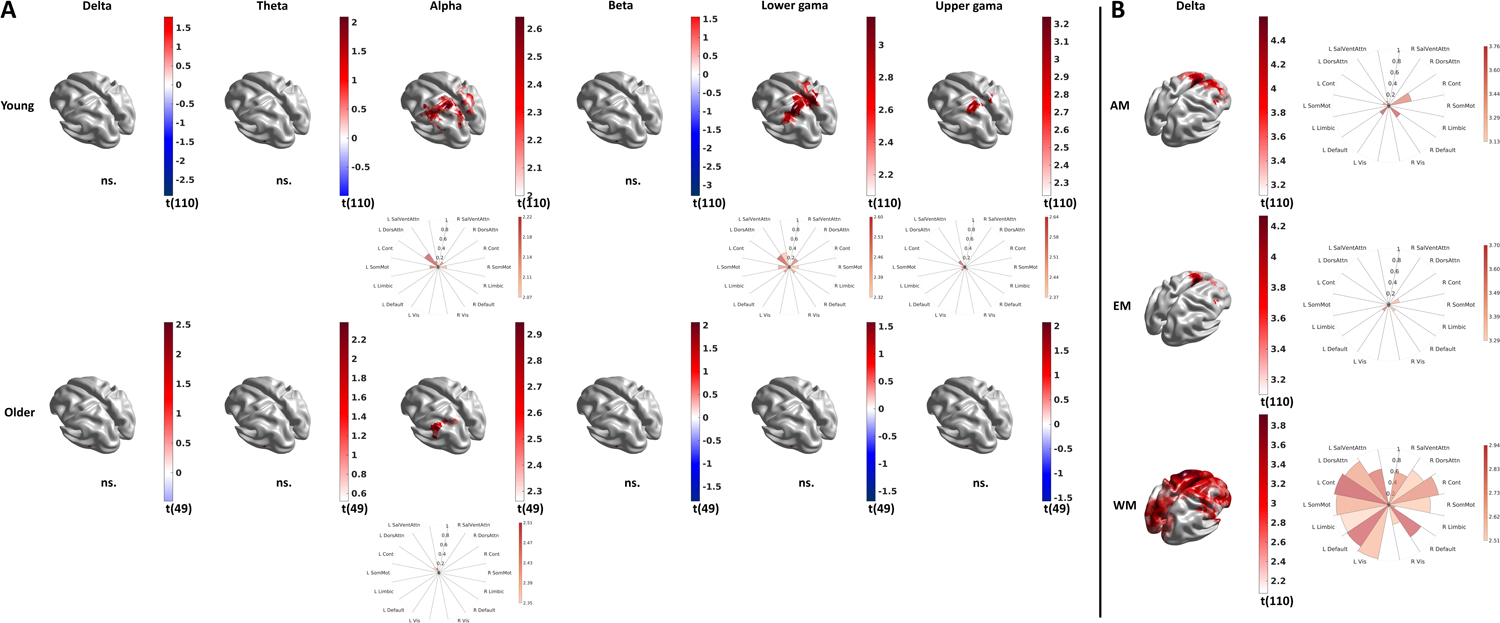
Neural correlates of memory. **(A)** The surface plots show brain areas where mean power (mECEO) in the individualised bands has a significant relationship with working memory performance of the “younger” (upper row) and the “older” (lower row) participants. **(B)** The surface plots show brain areas where reactivity (EC/EO) in the individualised delta band has a significant relationship with associative (AM), episodic (EM), and working memory (WM) performance of the “younger” participants. “ns.” denotes cased with no significant effect. The polar plots visualise the detected effects averaged in the seven functionally annotated networks in both hemispheres. The colour corresponds to the effect size as measured with the t-statistics of the regression, while the size of the wedges corresponds to the proportion of the functional network involved.

**Table 2.**
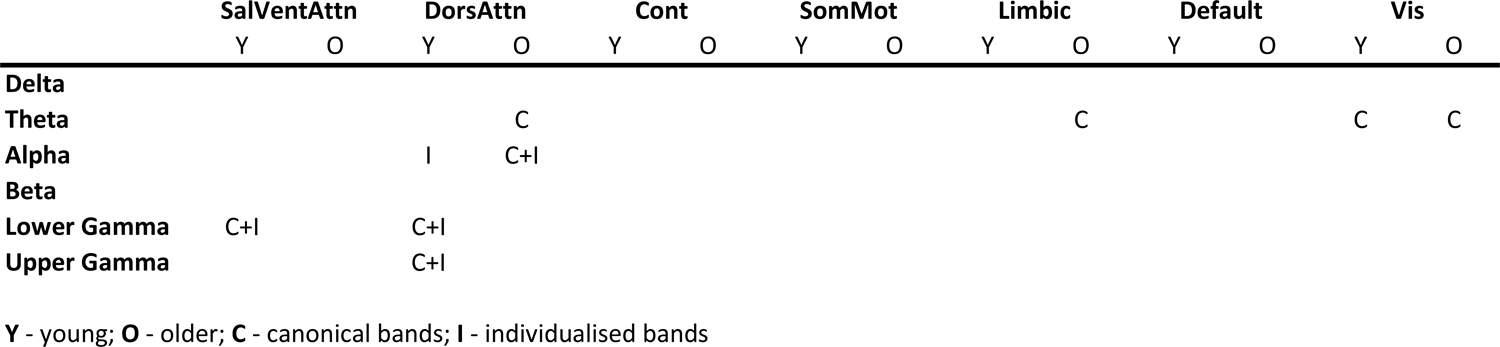
The effect band individualisation on the neural correlates of working memory. The table summarises in which functionally annotated networks we detected significant relationship between mean power (mECEO) and working memory performance of the “young” (Y) and the “older” (O) participants by using canonical (C) and individualised (I) bands.

When performing the same analyses for the canonical frequency bands we observed in addition a positive relationship between mEOEC theta power and working memory performance for both young and older adults (**Fig. S3** and **Table 2**). Band individualisation eliminated this relationship and improved the sensitivity to detect the abovementioned relationship in the alpha range.

Analyses of the relationship between memory performance and power reactivity estimates showed significantly positive relationships across all memory domains, but only for delta frequency and in the young adults’ group (**Fig. 6B**). This relationship was observed in the right Control and the bilateral Default networks in all three memory functions. For working memory there was a widely distributed correlation with delta reactivity. Band individualisation improved sensitivity in general and led to more significant test statistics and somewhat larger extent of the engagement of the various networks (**Fig. S3B**).

### 3.3. Age-related changes in connectivity in mEOEC and EO/EC reactivity and relation to memory performance

#### 3.3.1. Band individualisation effects on connectivity

Band individualisation affected connectivity estimates more drastically than power estimates. It eliminated all age-related and some of the WM-related variation in cross-frequency connectivity. For consistency and simplicity, in the rest of the paper we will only discuss cases where age- and memory-related findings were present for canonical and individualised bands.

#### 3.3.2. Age-related changes in connectivity

Aging lead to a widespread reduction in mEOEC delta connectivity, with stronger emphasis on the cross-hemispheric connections and a focal increase in the reactivity in lower gamma connectivity between the Control and Dorsal and Ventral Attention networks (**Fig. 7**). Band individualisation strongly increased the sensitivity (in the delta band) revealing a more distributed reduction in the connectivity, especially between the hemispheres (102 intra vs 266 cross connections). The involved connections form a network with the right Visual network (41% of the connections) as the main hub.

**Figure 7.**
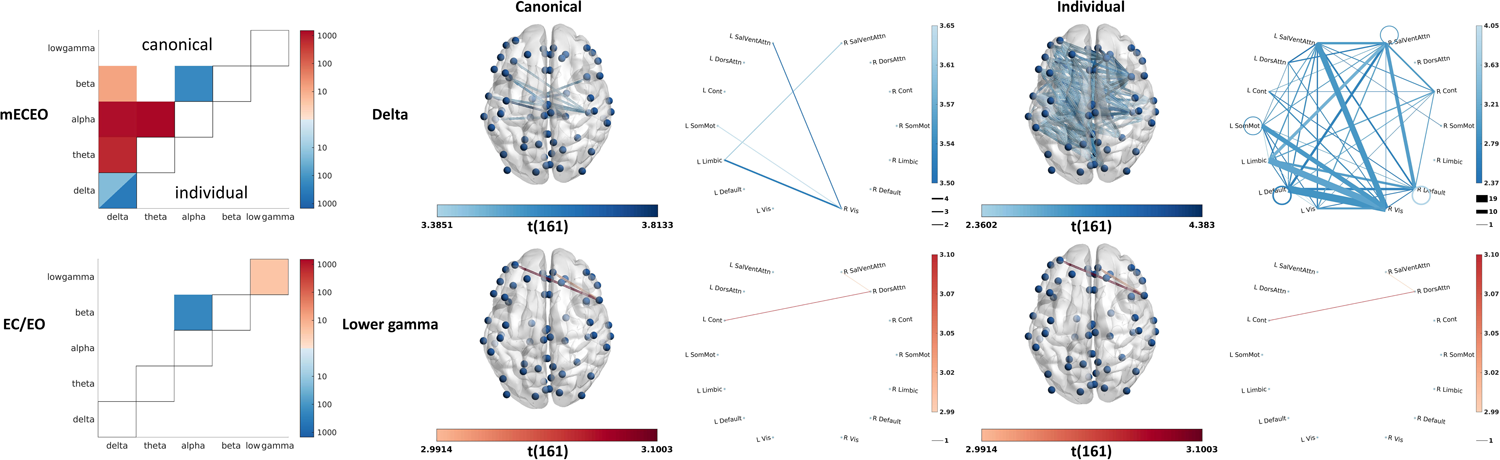
The effect of age and band individualisation on the combined connectivity measures. The matrix plots summarise the number of connections with significant age-related changes as detected by using canonical (upper triangle) and individualised (lower triangle) bands. Red and blue colours correspond to positive and negative effect, respectively. Within-frequency connectivity is visualised in the diagonal region, while cross-frequency connectivity is visualised in the off-diagonal regions. For the latter, only lower-to-higher frequency connectivity was estimated. For mean connectivity (mECEO), only the delta band showed significant age-related reduction also after band-individualisation, which is further visualised on the surface and the network plots.

#### 3.3.3. Connectivity correlates of memory

Analyses of the relationship between memory performance and the reactivity of connectivity revealed the involvement of several networks in working memory, however, only in the young group (**Fig. 8**). The reactivity of the theta-alpha, alpha-lower gamma, and beta-lower gamma connectivity across several networks showed to be increased in those with better working memory. The theta-alpha connections were rather intra-hemispheric (38 intra vs 22 cross) with right-sided dominance (9 left vs 29 right). The connectivity pattern formed a network with the right Ventral Attention (38% of the connections) and Visual (48% of the connections) networks as the main modulating hubs (62% and 93% of their connections are outputs, respectively). The alpha-lower gamma connections, however, were rather cross-hemispheric (42 intra vs 70 cross) with some within-hemispheric connections particularly in the left hemisphere (33 left vs 9 right). The connections for alpha-lower gamma formed a network with the left Visual (63% of the connections) network as the main receptive hub (80% of its connections are input).

**Figure 8.**
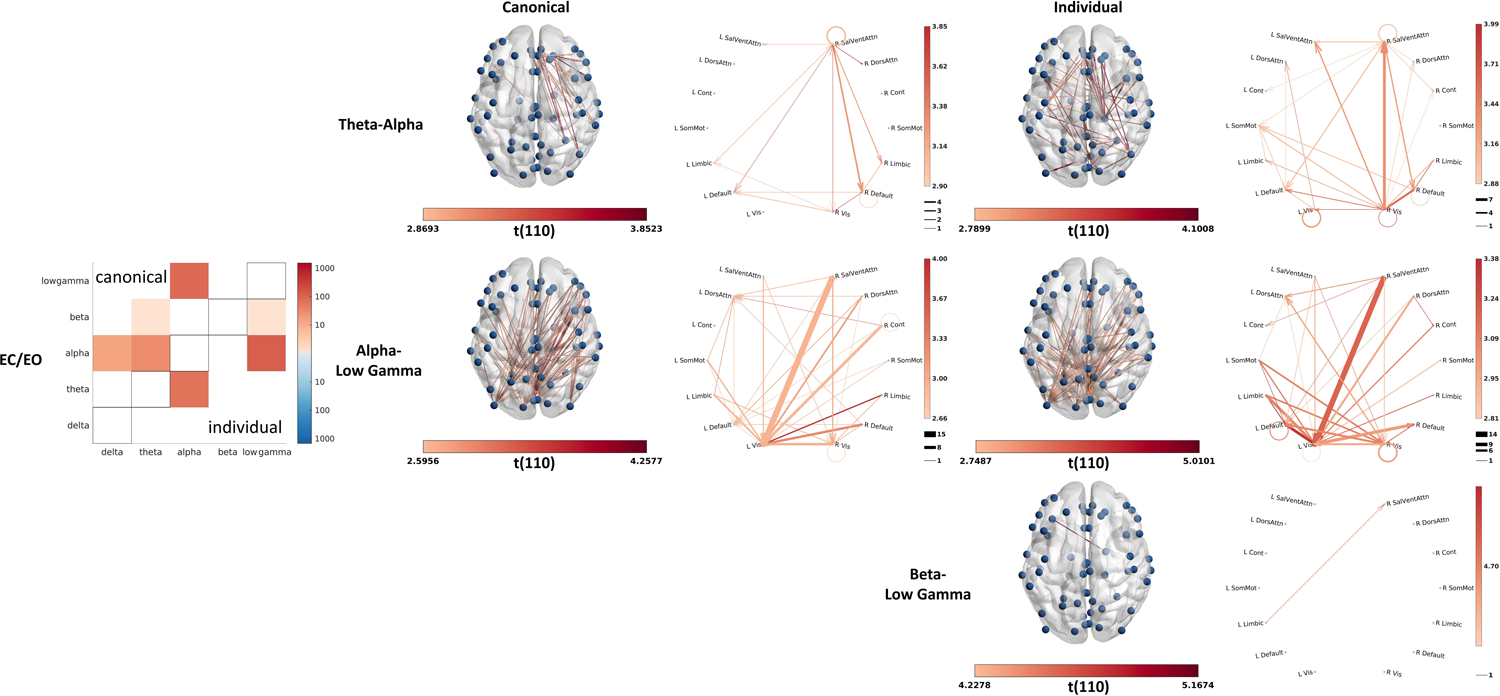
Connectivity correlates of working memory. The matrix plots summarise the number of connections with significant relationship with working memory performance of the “young” participants as detected by using canonical (upper triangle) and individualised (lower triangle) bands. Red and blue colours correspond to positive and negative effect, respectively. Within-frequency connectivity (none detected) is visualised in the diagonal region, while cross-frequency connectivity is visualised in the off-diagonal regions. For the latter, only lower-to-higher frequency connectivity was estimated. Only reactivity (EC/EO) in theta-alpha, alpha-lower gamma, and beta-lower gamma connectivity showed significant positive relationship with working memory performance of “young” participants also after band-individualisation, which are further visualised on the surface and the network plots. The cross-frequency connectivity is directed (lower to higher frequency), and the direction of the connectivity is marked with arrowheads.

## 4. Discussion

In this study, we used the open-access LEMON dataset to investigate electrophysiological markers of healthy ageing and memory function. Our study confirmed the impact of individually defined frequency bands on electrophysiological neuromarkers of ageing and cognitive function. Moreover, we also identified several resting state electrophysiological correlates of memory function, particularly when focusing on dynamic, i.e. reactivity estimates of power and functional connectivity. Notably, our findings are presented using functionally annotated brain networks to improve interpretability, we deploy graphs to perform cluster correction for functional connectivity and we provide a sharable and reproducible pipeline for electrophysiological data analysis.

### 4.1. Methodological implications

A robust finding in the literature is a slowing in alpha frequency with age (Cesnaite et al., 2023; Klimesch, 1999; Tröndle et al., 2023). The same observation was replicated here for the eyes closed condition where the alpha amplitude is naturally larger, and matching the condition typically investigated in previous ageing studies (Babiloni et al., 2006; Breslau et al., 1989; Polich, 1997; Tröndle et al., 2023; Vysata et al., 2012). This age-related shift in peak alpha frequency has important methodological implications, specifically that canonical alpha bands are not appropriate for investigating age-related changes in alpha power (Tröndle et al., 2023) or connectivity (Clark et al., 2004; Jabès et al., 2021; Knyazeva et al., 2018; Lodder and van Putten, 2011; Peltz et al., 2010). Our study confirms these findings and further suggests the importance of considering between-participant and -condition variability when adjusting bands. This band individualisation has a moderate-to-strong effect on potential neuromarkers. It moderately reduced delta and theta and strongly increased alpha and beta power (Fig. 4). Despite applying the individual band boundaries uniformly across the channels for each participant, we observed some topographic heterogeneity, especially for the alpha and beta bands, with a stronger effect in the posterior areas. More importantly, the effect of band individualisation on the reactivity indicates that EC and EO conditions are affected differently. This difference is negligible in the delta and theta bands (<3%) and moderate in the alpha (8-18%) and beta bands (5-40%). The topographic heterogeneity of the effect further supports an interaction between location and condition. These results underpin the importance of condition-specific band individualisation, which influences the sensitivity of identifying neural correlates of ageing and cognitive performance.

Since individual peaks were reliably detected only for the alpha and beta bands, corresponding differences, i.e., age-related increase in beta power (Fig. 5 vs Fig. S2) and neural correlates of working-memory in the alpha power (Fig. 6 vs Fig. S3), are not surprising. However, the downward shift of the alpha band can also affect the neighbouring theta band, which explains the increase in sensitivity of detecting age-related reduction in theta reactivity (Fig. 5 vs Fig. S2). The effect of band individualisation on connectivity is even more drastic. It eliminated most of the findings, while enabling greater sensitivity in detecting age-related increase in within-delta connectivity and some of the neural correlates of working memory in the reactivity of cross-frequency connectivity (Fig. 7-8).

The visual comparison of the magnitude of the effect of band individualisation on power and connectivity suggests that power measures are more robust and reliable because there is less difference between the two sets of findings. Connectivity, however, seems more sensitive to methodological choice; therefore, the construction of any workflow analysing connectivity should be well-justified. This also implies that findings on connectivity-based neuromarkers are more likely to change with the advancement of the field. Because methodological flexibility can lead to inconsistent findings and a lack of reproducibility (Lejko et al., 2020; Mahjoory et al., 2017) we used a reproducible and shareable pipeline (Automatic Analysis) to build a transparent workflow.

### 4.2. Confirmatory findings on age-related electrophysiological changes

The pace of reduction in alpha frequency with age (0.5 Hz from 20- to 80-year-old) was between what has been previously reported (0.45 Hz (Tröndle et al., 2023) to 1 Hz (Donoghue et al., 2020; Knyazeva et al., 2018). This slowing was observed throughout the scalp, especially in the midfrontal and occipital regions with larger effect on the right hemisphere, similarly to Tröndle et al. (Tröndle et al., 2023). Slowing in alpha frequency might reflect changes on the level of neurotransmission and excitation-inhibition ratios, as well as a decrease in axonal conduction velocity (Dustman et al., 1993; Hong and Rebec, 2012). A more controversial finding in ageing research pertains to changes in alpha power, with previous findings indicating either a decline with age (Babiloni et al., 2006; Barry and De Blasio, 2017; Kumral et al., 2020), or no age-related alterations (Sahoo et al., 2020) particularly when controlling for the reduction in alpha peak frequency or non-oscillatory 1/f slope of the EEG (Caplan et al., 2015; Cesnaite et al., 2023). Our findings agree with the later evidence of no changes in alpha power when defining individualised bands centred on individual alpha frequency.

We observed a reduction in delta and theta and an increase in beta power (mEOEC), similarly to the first study analysing this dataset with slightly different methodology (Kumral et al., 2020). The topography of age-related changes in mean power was somewhat similar to Kumral et al., 2020. However, there were also some notable differences, potentially due to a different estimation of power and our use of data acquired in both the EO and EC conditions. In contrast with Kumral et al., we found age-related reduction of mean theta power and reactivity in the occipital regions corresponding to the visual network and no changes in the midline regions. Age-related changes in theta power are however controversial. Previous studies have shown decreases (Vlahou et al., 2014), increases (Babiloni et al., 2006; Ishii et al., 2018; Klass and Brenner, 1995) or no changes (Hong and Rebec, 2012) with age. The observed decrease in delta and theta power with age in our study might however be explained by the now well-replicated flattening of the 1/f slope with age, with higher impact on lower frequencies (Dave et al., 2018; Lodder and van Putten, 2011; Voytek et al., 2015). Perhaps less controversial is our observation of age-related increases in beta power, which agrees with previous literature showing higher resting-state beta activity in older adults (Gómez et al., 2013; Heinrichs-Graham et al., 2018; Heinrichs-Graham and Wilson, 2016; Hübner et al., 2018; Koyama et al., 1997; Veldhuizen et al., 1993). The topography of our findings matches previous studies showing higher spontaneous beta power in frontal and parietal regions that form part of the Somatomotor and Salience networks. In particular, the observation of increased power in the Somatomotor network matches previous whole-brain analysis indicating motor regions as those with higher differences in beta power between young and older cohorts (Heinrichs-Graham and Wilson, 2016). Changes in beta power at rest are thought to index age-related changes in GABAergic inhibition (Inamoto et al., 2023; Rossiter et al., 2014).

In regards to age-related changes in reactivity we observed a decrease in reactivity across all frequency bands with the exception of the alpha band. In accordance with Barry and De Blasio (Barry and De Blasio, 2017), we also detected an anterior-posterior gradient in the spatial distribution of age-related changes in the reactivity across several bands. We complement these findings by showing a reduction in gamma reactivity, which was not previously investigated in this dataset but is consistent with similar findings in Jabes et al. (Jabès et al., 2021). Moreover, our findings allow us to observe finer spatial patterns at functional network level (see below) that were not captured by previous studies with lower spatial resolution.

The analysis of the relationship between connectivity and age revealed several age-related changes. While most age-related changes in mEOEC power were not significant when using individualised bands, we observed a robust age-related decrease in connectivity in the delta band across hemispheres, with most changes affecting the connections of the right visual network. Comparison of resting state EEG of Alzheimer’s disease patients and healthy controls led to similar findings suggesting a decline in interhemispheric communication or compensation mechanisms (Hata et al., 2016).

### 4.3. New findings on age-related electrophysiological changes and memory function

With our more comprehensive analysis, we were also able to complement existing findings and contribute to the debate on the relationship between EEG oscillations and cognitive ageing. We propose that our assessment of the effect of the condition-specific individualisation adds additional weight to our findings. In particular, we found increased sensitivity in detecting age-related changes in the beta power, the theta reactivity and the delta connectivity and the neural correlates of working-memory in the alpha power and the theta-alpha and the alpha-lower gamma connectivity.

Several studies, however, found a link between the alpha power and cognitive performance or decline (Babiloni et al., 2022; Fonseca et al., 2011; Van Der Hiele et al., 2008), which is also consistent with our finding of a positive correlation between alpha (and gamma) power and working memory performance. These indicate that alpha power can be considered a reliable and specific neuromarker of working memory performance irrespective of age.

However, our investigation of the relationship between functional connectivity and memory performance only showed consistent findings in the younger group. In particular, we found that better working memory performance correlated with rest reactivity of theta-alpha connectivity within and between frontal and parietal regions. This pattern closely resembles changes elicited by working-memory tasks (Akiyama et al., 2017). Moreover, the reactivity of both theta-alpha and alpha-gamma connectivity have been reported to correlate with attention, processing speed, and executive functions (Siebenhühner et al., 2020), key components of working memory. Our findings also resolve some of the inconsistency around the role of delta oscillations in cognition (Trammell et al., 2017) by showing that it reflects domain-independent dynamics (i.e., reactivity) rather than static components of memory. The lack of significant correlates of connectivity and memory performance in older adults might be partly explained by the lower sample size in comparison with the effect size in this age group; however, the literature on the relationship between rs-EEG connectivity and memory is scarce. Hata and coworkers, for example, were able to demonstrate the reduced connectivity within the delta band as a potential neuromarker of dementia rather than healthy memory function in a sample including 28 “probable AD patients” and 30 healthy controls (Hata et al., 2016).

#### 4.3.1. Functionally annotated electrophysiology

Our electrophysiological findings confirmed that ageing affects the activity (power) and reactivity of multiple networks involved in perception and cognition, such as the Control, Dorsal Attention, Visual, Default networks (Fig. 5). The functional annotation draws a richer picture of and provides contextual support and mechanistic understanding for the neural correlates of memory performance, indicating the primary roles of Dorsal and Ventral Attention networks in working memory (Fig. 6a) and the primary role of Control networks in associative and episodic memory (Fig. 6b). The left hemispheric lateralisation of the neural correlates of working memory (Fig. 6a) emphasizes the verbal components (Nagel et al., 2013; Ray et al., 2008) of the working memory task used during the assessment. Our findings on the role of cross-frequency connectivity in working memory further confirms the sensitivity of neuromarkers related to dynamic (i.e., reactivity) rather than static neural features and revealed some fundamental differences in the engagement of the lower-(theta-alpha) and higher-(alpha-gamma) frequency networks in working memory. The lower-frequency network predominantly engages the right hemisphere via shorter paths and is driven by the Visual and Ventral Attention network (Fig. 8, first row) suggesting integration of information implemented via perception driven (i.e., bottom-up) and salience and subsequent attention-driven (i.e., top-down) processes (Akiyama et al., 2017). The higher-frequency network, on the other hand, is characterised by cross-hemispheric communication via longer paths modulated by the Visual network (Fig. 8, second row) suggesting the dominance of top-down processes.

Investigating electrophysiological neuromarkers at source level allows one to identify their correspondence and functional annotation with fMRI findings even without synchronous recording. In addition to improving the interpretability of the EEG findings, this approach also allows to put them into a wider multimodal context.

### 4.4. Limitations

While the LEMON dataset constitutes a powerful resource to study electrophysiological markers of healthy ageing, it currently has some inherent limitations. The first is the absence of participants’ ages, as the dataset contains instead nine pre-defined age groups. Such arrangement reduces the sensitivity to detect age-related changes because it does not allow the full modelling of age-related variability. The second is the overrepresentation of younger participants (<=35 year-old) in the sample. This was further aggravated by the rejection of samples with artifacts, which was more common in the older group. As a result, our analyses have more power to detect effects in the “young” than in the “older” group; therefore, the lack of an effect in the “older” group cannot be unambiguously interpreted. Thirdly, we omitted the investigation of the effect of sex due to the unbalanced representation of females and males across the age groups and the whole sample. Finally, information about the time of the day in which acquisitions took place was not available and circadian and ultradian rhythms can impact spectral characteristics of EEG signals (Lehnertz et al., 2021). These limitations might be addressed by the addition of further information and extensions to the LEMON dataset or combining it with other openly available datasets. The analyses of these datasets will benefit from reproducible pipelines, such as the one we produced for this study.

An additional limitation of our study is that power estimates included both periodic and aperiodic components. While we did account for individual differences when defining frequency bands, a recent study emphasized the importance of decomposing periodic and aperiodic components when investigating the age effect on alpha power during the EC conditions, albeit mentioning that the contribution of the two components to cognitive decline is still unclear (Tröndle et al., 2023). Future studies could investigate how accounting for both individualised bands and aperiodic components contribute to changes in functional connectivity patterns within and between frequency bands and their relation to cognitive performance.

Finally, the statistical framework as implemented in Fieldtrip does not support multiple regression. To partially overcome this limitation, we split the participants into “young” and “older” groups, however, it does not allow a full separation of the age-effect from the correlates of memory functions, somewhat reducing the sensitivity to detect them. The LIMO toolbox allows modelling multiple variabilities in M/EEG data by using hierarchical linear modelling (Pernet et al., 2011). Future developments in our pipeline or Fieldtrip might address this issue and allow the full integration of LIMO.

## 5. Conclusion

Despite the recent developments in neuroimaging, which allow sophisticated analyses on larger datasets to extract psychobiologically relevant features to be conducted, we still have limited success in identifying reliable neuromarkers of age-related cognitive decline. The present paper synthesizes best practices and implements methodological considerations to provide an integrative insight into the electrophysiological correlates of age-related cognitive decline using functionally annotated brain networks. Our analyses confirmed the most established neuromarkers of cognitive ageing, while it also clarified the role of delta and alpha oscillations in memory performance and their changes in ageing. We also highlighted the importance of considering the variation in band specification across individuals and conditions and demonstrated the higher sensitivity of dynamic rather than static electrophysiological neuromarkers in ageing and memory.

## 6. Author contributions

**Tibor Auer**: Conceptualisation, Data curation, Methodology, Software, Formal analysis, Validation, Writing - Original Draft, Writing - Review & Editing, Visualization, Supervision; **Robin Goldthorpe**: Conceptualisation, Data curation, Writing - Original Draft; **Robert Peach**: Conceptualization, Writing - Review & Editing; **Henry Hebron:** Methodology, Software, Formal analysis; **Ines Violante**: Conceptualisation, Supervision, Funding acquisition, Project administration, Writing - Original Draft, Writing - Review & Editing

## 7. Funding

This work was supported by the Biotechnology and Biological Sciences Research Council, London [BB/S008314].

## 8. Declarations of interest

None

## 11. Supplemental materials

**Supplemental figure 1. Data processing and analysis workflow as implemented in Automatic Analysis (aa)**

The figure represents the data processing and analysis steps and the data flow between them. The steps, implemented as aa modules (aamod_*), are visualised as text in a box, while the data, managed in aa streams, are visualised as text on arrows indicating direction of the dataflow. The preprocessing steps are common for the canonical and individualised bands. Analysis steps using canonical bands are marked with green, while analysis steps using individualised bands are marked with turquoise.

**Supplemental figure 2. Age-related changes in the combined neural measures**

The surface plots demonstrate the significant localised effect of ageing, measured as t-statistics of the regression, on the combined neural measures of mean power (mECEO) and reactivity (EC/EO) in the canonical bands. “ns.” denotes cased with no significant effect. The polar plots visualise the detected effects averaged in the seven functionally annotated networks in both hemispheres. The colour corresponds to the effect size as measured with the t-statistics of the regression, while the size of the wedges corresponds to the proportion of the functional network involved.

**Supplemental figure 3. Neural correlates of memory**

**(A)** The surface plots show brain areas where mean power (mECEO) in the canonical bands has a significant relationship with working memory performance of the “younger” (upper row) and the “older” (lower row) participants. **(B)** The surface plots show brain areas where reactivity (EC/EO) in the canonical delta band has a significant relationship with associative (AM), episodic (EM), and working memory (WM) performance of the “younger” participants.

“ns.” denotes cased with no significant effect. The polar plots visualise the detected effects averaged in the seven functionally annotated networks in both hemispheres. The colour corresponds to the effect size as measured with the t-statistics of the regression, while the size of the wedges corresponds to the proportion of the functional network involved.

**Supplemental table 1. Participant demographics**

The table summarises the number of participants divided into sexes and the nine age groups in the final sample after data cleaning.

**Supplemental table 2. Memory performance**

The table summarises the associative (AM), episodic (EM), and working memory (WM) performance of the “young” and the “older” participants.

## Notes

### Competing Interest Statement

The authors have declared no competing interest.

### Summary of Updates

Amend "Author contribution"

## References

Akiyama, M., Tero, A., Kawasaki, M., Nishiura, Y., Yamaguchi, Y., 2017. Theta-alpha EEG phase distributions in the frontal area for dissociation of visual and auditory working memory. Sci. Rep. 7, 1–11. 10.1038/srep42776

Babayan, A., Erbey, M., Kumral, D., Reinelt, J.D., Reiter, A.M.F., Röbbig, J., Lina Schaare, H., Uhlig, M., Anwander, A., Bazin, P.L., Horstmann, A., Lampe, L., Nikulin, V. V., Okon-Singer, H., Preusser, S., Pampel, A., Rohr, C.S., Sacher, J., Thöne-Otto, A., Trapp, S., Nierhaus, T., Altmann, D., Arelin, K., Blöchl, M., Bongartz, E., Breig, P., Cesnaite, E., Chen, S., Cozatl, R., Czerwonatis, S., Dambrauskaite, G., Dreyer, M., Enders, J., Engelhardt, M., Fischer, M.M., Forschack, N., Golchert, J., Golz, L., Guran, C.A., Hedrich, S., Hentschel, N., Hoffmann, D.I., Huntenburg, J.M., Jost, R., Kosatschek, A., Kunzendorf, S., Lammers, H., Lauckner, M.E., Mahjoory, K., Kanaan, A.S., Mendes, N., Menger, R., Morino, E., Näthe, K., Neubauer, J., Noyan, H., Oligschläger, S., Panczyszyn-Trzewik, P., Poehlchen, D., Putzke, N., Roski, S., Schaller, M.C., Schieferbein, A., Schlaak, B., Schmidt, R., Gorgolewski, K.J., Schmidt, H.M., Schrimpf, A., Stasch, S., Voss, M., Wiedemann, A., Margulies, D.S., Gaebler, M., Villringer, A., 2019. Data descriptor: A mind-brain-body dataset of MRI, EEG, cognition, emotion, and peripheral physiology in young and old adults. Sci. Data 6, 1–21. 10.1038/sdata.2018.308

Babiloni, C., Binetti, G., Cassarino, A., Dal Forno, G., Del Percio, C., Ferreri, F., Ferri, R., Frisoni, G., Galderisi, S., Hirata, K., Lanuzza, B., Miniussi, C., Mucci, A., Nobili, F., Rodriguez, G., Romani, G.L., Rossini, P.M., 2006. Sources of cortical rhythms in adults during physiological aging: A multicentric EEG study. Hum. Brain Mapp. 27, 162–172. 10.1002/HBM.20175

Babiloni, C., Lorenzo, I., Lizio, R., Lopez, S., Tucci, F., Ferri, R., Soricelli, A., Nobili, F., Arnaldi, D., Famà, F., Buttinelli, C., Giubilei, F., Cipollini, V., Onofrj, M., Stocchi, F., Vacca, L., Fuhr, P., Gschwandtner, U., Ransmayr, G., Aarsland, D., Parnetti, L., Marizzoni, M., D’Antonio, F., De Lena, C., Güntekin, B., Yıldırım, E., Hanoğlu, L., Yener, G., Gündüz, D.H., Taylor, J.P., Schumacher, J., McKeith, I., Frisoni, G.B., De Pandis, M.F., Bonanni, L., Percio, C. Del, Noce, G., 2022. Reactivity of posterior cortical electroencephalographic alpha rhythms during eyes opening in cognitively intact older adults and patients with dementia due to Alzheimer’s and Lewy body diseases. Neurobiol. Aging 115, 88–108. 10.1016/j.neurobiolaging.2022.04.001

Barry, R.J., De Blasio, F.M., 2017. EEG differences between eyes-closed and eyes-open resting remain in healthy ageing. Biol. Psychol. 129, 293–304. 10.1016/j.biopsycho.2017.09.010

Başar, E., 2013. A review of gamma oscillations in healthy subjects and in cognitive impairment. Int. J. Psychophysiol. 10.1016/j.ijpsycho.2013.07.005

Bellato, A., Arora, I., Kochhar, P., Hollis, C., Groom, M.J., 2020. Atypical electrophysiological indices of eyes-open and eyes-closed resting-state in children and adolescents with ADHD and autism. Brain Sci. 10, 272. 10.3390/brainsci10050272

Belleville, S., Bherer, L., 2012. Biomarkers of Cognitive Training Effects in Aging. Curr. Transl. Geriatr. Exp. Gerontol. Rep. 1, 104–110. 10.1007/s13670-012-0014-5

Breslau, J., Starr, A., Sicotte, N., Higa, J., Buchsbaum, M.S., 1989. Topographic EEG changes with normal aging and SDAT. Electroencephalogr. Clin. Neurophysiol. 72, 281–289. 10.1016/0013-4694(89)90063-1

Brookes, M.J., Woolrich, M., Luckhoo, H., Price, D., Hale, J.R., Stephenson, M.C., Barnes, G.R., Smith, S.M., Morris, P.G., 2011. Investigating the electrophysiological basis of resting state networks using magnetoencephalography. Proc. Natl. Acad. Sci. U. S. A. 108, 16783–16788. 10.1073/pnas.1112685108

Caplan, J.B., Bottomley, M., Kang, P., Dixon, R.A., 2015. Distinguishing rhythmic from non-rhythmic brain activity during rest in healthy neurocognitive aging. Neuroimage 112, 341–352. 10.1016/j.neuroimage.2015.03.001

Cesnaite, E., Steinfath, P., Jamshidi Idaji, M., Stephani, T., Kumral, D., Haufe, S., Sander, C., Hensch, T., Hegerl, U., Riedel-Heller, S., Röhr, S., Schroeter, M.L., Witte, A., Villringer, A., Nikulin, V. V., 2023. Alterations in rhythmic and non-rhythmic resting-state EEG activity and their link to cognition in older age. Neuroimage 268, 119810. 10.1016/J.NEUROIMAGE.2022.119810

Chang, C.Y., Hsu, S.H., Pion-Tonachini, L., Jung, T.P., 2018. Evaluation of Artifact Subspace Reconstruction for Automatic EEG Artifact Removal, in: Proceedings of the Annual International Conference of the IEEE Engineering in Medicine and Biology Society, EMBS. Institute of Electrical and Electronics Engineers Inc., pp. 1242–1245. 10.1109/EMBC.2018.8512547

Choi, K.M., Kim, J.Y., Kim, Y.W., Han, J.W., Im, C.H., Lee, S.H., 2021. Comparative analysis of default mode networks in major psychiatric disorders using resting-state EEG. Sci. Rep. 11, 1–10. 10.1038/s41598-021-00975-3

Chow, R., Rabi, R., Paracha, S., Hasher, L., Anderson, N.D., Alain, C., 2022. Default Mode Network and Neural Phase Synchronization in Healthy Aging: A Resting State EEG Study. Neuroscience 485, 116–128. 10.1016/j.neuroscience.2022.01.008

Christensen, K., Doblhammer, G., Rau, R., Vaupel, J.W., 2009. Ageing populations: the challenges ahead. Lancet. 10.1016/S0140-6736(09)61460-4

Clark, C.R., Veltmeyer, M.D., Hamilton, R.J., Simms, E., Paul, R., Hermens, D., Gordon, E., 2004. Spontaneous alpha peak frequency predicts working memory performance across the age span. Int. J. Psychophysiol. 53, 1–9. 10.1016/J.IJPSYCHO.2003.12.011

Cohen, M.X., 2021. A data-driven method to identify frequency boundaries in multichannel electrophysiology data. J. Neurosci. Methods 347, 108949. 10.1016/j.jneumeth.2020.108949

Cole, J.H., Franke, K., 2017. Predicting Age Using Neuroimaging: Innovative Brain Ageing Biomarkers. Trends Neurosci. 10.1016/j.tins.2017.10.001

Collerton, J., Barrass, K., Bond, J., Eccles, M., Jagger, C., James, O., Martin-Ruiz, C., Robinson, L., Von Zglinicki, T., Kirkwood, T., 2007. The Newcastle 85+ study: Biological, clinical and psychosocial factors associated with healthy ageing: Study protocol. BMC Geriatr. 7. 10.1186/1471-2318-7-14

Cusack, R., Vicente-Grabovetsky, A., Mitchell, D.J., Wild, C.J., Auer, T., Linke, A.C., Peelle, J.E., 2015. Automatic analysis (aa): Efficient neuroimaging workflows and parallel processing using Matlab and XML. Front. Neuroinform. 8, 90. 10.3389/fninf.2014.00090

Custo, A., Van De Ville, D., Wells, W.M., Tomescu, M.I., Brunet, D., Michel, C.M., 2017. Electroencephalographic Resting-State Networks: Source Localization of Microstates. Brain Connect. 7, 671–682. 10.1089/brain.2016.0476

Dave, S., Brothers, T.A., Swaab, T.Y., 2018. 1/f neural noise and electrophysiological indices of contextual prediction in aging. Brain Res. 1691, 34–43. 10.1016/j.brainres.2018.04.007

Delorme, A., Makeig, S., 2004. EEGLAB: An open source toolbox for analysis of single-trial EEG dynamics including independent component analysis. J. Neurosci. Methods 134, 9–21. 10.1016/j.jneumeth.2003.10.009

Delorme, A., Palmer, J., Onton, J., Oostenveld, R., Makeig, S., 2012. Independent EEG sources are dipolar. PLoS One 7, 30135. 10.1371/journal.pone.0030135

Desikan, R.S., Ségonne, F., Fischl, B., Quinn, B.T., Dickerson, B.C., Blacker, D., Buckner, R.L., Dale, A.M., Maguire, R.P., Hyman, B.T., Albert, M.S., Killiany, R.J., 2006. An automated labeling system for subdividing the human cerebral cortex on MRI scans into gyral based regions of interest. Neuroimage 31, 968–980. 10.1016/j.neuroimage.2006.01.021

Dewiputri, W.I., Schweizer, R., Auer, T., 2021. Brain Networks Underlying Strategy Execution and Feedback Processing in an Efficient Functional Magnetic Resonance Imaging Neurofeedback Training Performed in a Parallel or a Serial Paradigm. Front. Hum. Neurosci. 15, 215. 10.3389/fnhum.2021.645048

Dickerson, B.C., Wolk, D.A., 2012. MRI cortical thickness biomarker predicts AD-like CSF and cognitive decline in normal adults. Neurology 78, 84–90. 10.1212/WNL.0b013e31823efc6c

Donoghue, T., Haller, M., Peterson, E.J., Varma, P., Sebastian, P., Gao, R., Noto, T., Lara, A.H., Wallis, J.D., Knight, R.T., Shestyuk, A., Voytek, B., 2020. Parameterizing neural power spectra into periodic and aperiodic components. Nat. Neurosci. 23, 1655–1665. 10.1038/s41593-020-00744-x

Dustman, R.E., Shearer, D.E., Emmerson, R.Y., 1993. EEG and event-related potentials in normal aging. Prog. Neurobiol. 10.1016/0301-0082(93)90005-D

Engels, M.M.A., Stam, C.J., van der Flier, W.M., Scheltens, P., de Waal, H., van Straaten, E.C.W., 2015. Declining functional connectivity and changing hub locations in Alzheimer’s disease: An EEG study. BMC Neurol. 15, 1–8. 10.1186/s12883-015-0400-7

Engemann, D.A., Kozynets, O., Sabbagh, D., Lemaitre, G., Varoquaux, G., Liem, F., Gramfort, A., 2020. Combining magnetoencephalography with magnetic resonance imaging enhances learning of surrogate-biomarkers. Elife 9, 1–33. 10.7554/eLife.54055

Fonseca, L.C., Tedrus, G.M.A.S., Fondello, M.A., Reis, I.N., Fontoura, D.S., 2011. EEG theta and alpha reactivity on opening the eyes in the diagnosis of Alzheimer’s disease. Clin. EEG Neurosci. 42, 185–189. 10.1177/155005941104200308

Foo, H., Thalamuthu, A., Jiang, J., Koch, F.C., Mather, K.A., Wen, W., Sachdev, P.S., 2021. Novel genetic variants associated with brain functional networks in 18,445 adults from the UK Biobank. Sci. Rep. 11, 1–12. 10.1038/s41598-021-94182-9

Fox, M.D., Raichle, M.E., 2007. Spontaneous fluctuations in brain activity observed with functional magnetic resonance imaging. Nat. Rev. Neurosci. 10.1038/nrn2201

Frömer, R., Maier, M., Abdel Rahman, R., 2018. Group-Level EEG-Processing Pipeline for Flexible Single Trial-Based Analyses Including Linear Mixed Models. Front. Neurosci. 12, 48. 10.3389/fnins.2018.00048

Funder, D.C., Ozer, D.J., 2019. Evaluating Effect Size in Psychological Research: Sense and Nonsense. Adv. Methods Pract. Psychol. Sci. 2, 156–168. 10.1177/2515245919847202

Gallen, C.L., D’Esposito, M., 2019. Brain Modularity: A Biomarker of Intervention-related Plasticity. Trends Cogn. Sci. 10.1016/j.tics.2019.01.014

Gómez, C., M Pérez-Macías, J., Poza, J., Fernández, A., Hornero, R., 2013. Spectral changes in spontaneous MEG activity across the lifespan. J. Neural Eng. 10, 066006. 10.1088/1741-2560/10/6/066006

Grady, C., 2012. The cognitive neuroscience of ageing. Nat. Rev. Neurosci. 10.1038/nrn3256

Hadia, N.U., Abdullah, N., Sentosa, I., 2016. An Easy Approach to Exploratory Factor Analysis: Marketing Perspective. J. Educ. Soc. Res. 6. 10.5901/jesr.2016.v6n1p215

Harary, F., Norman, R.Z., 1960. Some properties of line digraphs. Rend. del Circ. Mat. di Palermo 9, 161–168. 10.1007/BF02854581

Hata, M., Kazui, H., Tanaka, T., Ishii, R., Canuet, L., Pascual-Marqui, R.D., Aoki, Y., Ikeda, S., Kanemoto, H., Yoshiyama, K., Iwase, M., Takeda, M., 2016. Functional connectivity assessed by resting state EEG correlates with cognitive decline of Alzheimer’s disease - An eLORETA study. Clin. Neurophysiol. 127, 1269–1278. 10.1016/j.clinph.2015.10.030

Heinrichs-Graham, E., McDermott, T.J., Mills, M.S., Wiesman, A.I., Wang, Y.P., Stephen, J.M., Calhoun, V.D., Wilson, T.W., 2018. The lifespan trajectory of neural oscillatory activity in the motor system. Dev. Cogn. Neurosci. 30, 159–168. 10.1016/j.dcn.2018.02.013

Heinrichs-Graham, E., Wilson, T.W., 2016. Is an absolute level of cortical beta suppression required for proper movement? Magnetoencephalographic evidence from healthy aging. Neuroimage 134, 514–521. 10.1016/j.neuroimage.2016.04.032

Herrmann, C.S., Fründ, I., Lenz, D., 2010. Human gamma-band activity: A review on cognitive and behavioral correlates and network models. Neurosci. Biobehav. Rev. 10.1016/j.neubiorev.2009.09.001

Herrmann, C.S., Munk, M.H.J., Engel, A.K., 2004. Cognitive functions of gamma-band activity: Memory match and utilization. Trends Cogn. Sci. 8, 347–355. 10.1016/j.tics.2004.06.006

Hillebrand, A., Barnes, G.R., Bosboom, J.L., Berendse, H.W., Stam, C.J., 2012. Frequency-dependent functional connectivity within resting-state networks: An atlas-based MEG beamformer solution. Neuroimage 59, 3909–3921. 10.1016/j.neuroimage.2011.11.005

Hong, S.L., Rebec, G. V., 2012. A new perspective on behavioral inconsistency and neural noise in aging: Compensatory speeding of neural communication. Front. Aging Neurosci. 4, 33404. 10.3389/fnagi.2012.00027

Huang, Y., Parra, L.C., Haufe, S., 2016. The New York Head—A precise standardized volume conductor model for EEG source localization and tES targeting. Neuroimage 140, 150–162. 10.1016/j.neuroimage.2015.12.019

Hübner, L., Godde, B., Voelcker-Rehage, C., 2018. Older adults reveal enhanced task-related beta power decreases during a force modulation task. Behav. Brain Res. 345, 104–113. 10.1016/j.bbr.2018.02.028

Inamoto, T., Ueda, M., Ueno, K., Shiroma, C., Morita, R., Naito, Y., Ishii, R., 2023. Motor-Related Mu/Beta Rhythm in Older Adults: A Comprehensive Review. Brain Sci. 10.3390/brainsci13050751

Ishii, R., Canuet, L., Aoki, Y., Hata, M., Iwase, M., Ikeda, S., Nishida, K., Ikeda, M., 2018. Healthy and Pathological Brain Aging: From the Perspective of Oscillations, Functional Connectivity, and Signal Complexity. Neuropsychobiology. 10.1159/000486870

Jabès, A., Klencklen, G., Ruggeri, P., Antonietti, J.P., Banta Lavenex, P., Lavenex, P., 2021. Age-Related Differences in Resting-State EEG and Allocentric Spatial Working Memory Performance. Front. Aging Neurosci. 13, 715. 10.3389/fnagi.2021.704362

Jack, C.R., Wiste, H.J., Weigand, S.D., Therneau, T.M., Lowe, V.J., Knopman, D.S., Gunter, J.L., Senjem, M.L., Jones, D.T., Kantarci, K., Machulda, M.M., Mielke, M.M., Roberts, R.O., Vemuri, P., Reyes, D.A., Petersen, R.C., 2017. Defining imaging biomarker cut points for brain aging and Alzheimer’s disease. Alzheimer’s Dement. 13, 205–216. 10.1016/j.jalz.2016.08.005

James, L.E., Fogler, K.A., Tauber, S.K., 2008. Recognition Memory Measures Yield Disproportionate Effects of Aging on Learning Face-Name Associations. Psychol. Aging 23, 657–664. 10.1037/a0013008

Javaid, H., Kumarnsit, E., Chatpun, S., 2022. Age-Related Alterations in EEG Network Connectivity in Healthy Aging. Brain Sci. 2022, Vol. 12, Page 218 12, 218. 10.3390/BRAINSCI12020218

Jokisch, D., Jensen, O., 2007. Modulation of gamma and alpha activity during a working memory task engaging the dorsal or ventral stream. J. Neurosci. 27, 3244–3251. 10.1523/JNEUROSCI.5399-06.2007

Jonker, C., Jonker, M.I., Schmand, B., 2000. Are memory complaints predictive for dementia? A review of clinical and population-based studies. Int. J. Geriatr. Psychiatry. 10.1002/1099-1166(200011)15:11<983::AID-GPS238>3.0.CO;2-5

Klass, D.W., Brenner, R.P., 1995. Electroencephalography of the Elderly. J. Clin. Neurophysiol. 12.

Klimesch, W., 1999. EEG alpha and theta oscillations reflect cognitive and memory performance: A review and analysis. Brain Res. Rev. 10.1016/S0165-0173(98)00056-3

Knyazeva, M.G., Barzegaran, E., Vildavski, V.Y., Demonet, J.F., 2018. Aging of human alpha rhythm. Neurobiol. Aging 69, 261–273. 10.1016/j.neurobiolaging.2018.05.018

Kosciessa, J.Q., Grandy, T.H., Garrett, D.D., Werkle-Bergner, M., 2020. Single-trial characterization of neural rhythms: Potential and challenges. Neuroimage 206, 116331. 10.1016/j.neuroimage.2019.116331

Koyama, K., Hirasawa, H., Okubo, Y., Karasawa, A., 1997. Quantitative EEG Correlates of Normal Aging in the Elderly. Clin. EEG Neurosci. 28, 160–165. 10.1177/155005949702800308

Kumral, D., Şansal, F., Cesnaite, E., Mahjoory, K., Al, E., Gaebler, M., Nikulin, V. V., Villringer, A., 2020. BOLD and EEG signal variability at rest differently relate to aging in the human brain. Neuroimage 207, 116373. 10.1016/j.neuroimage.2019.116373

Lehnertz, K., Rings, T., Bröhl, T., 2021. Time in Brain: How Biological Rhythms Impact on EEG Signals and on EEG-Derived Brain Networks. Front. Netw. Physiol. 1, 755016. 10.3389/fnetp.2021.755016

Lejko, N., Larabi, D.I., Herrmann, C.S., Aleman, A., Ćurčić-Blake, B., 2020. Alpha Power and Functional Connectivity in Cognitive Decline: A Systematic Review and Meta-Analysis. J. Alzheimer’s Dis. 10.3233/JAD-200962

Lodder, S.S., van Putten, M.J.A.M., 2011. Automated EEG analysis: Characterizing the posterior dominant rhythm. J. Neurosci. Methods 200, 86–93. 10.1016/j.jneumeth.2011.06.008

Mahjoory, K., Nikulin, V. V., Botrel, L., Linkenkaer-Hansen, K., Fato, M.M., Haufe, S., 2017. Consistency of EEG source localization and connectivity estimates. Neuroimage 152, 590–601. 10.1016/j.neuroimage.2017.02.076

Majerus, S., Péters, F., Bouffier, M., Cowan, N., Phillips, C., 2018. The dorsal attention network reflects both encoding load and top-down control during working memory. J. Cogn. Neurosci. 30, 144–159. 10.1162/jocn_a_01195

Maris, E., Oostenveld, R., 2007. Nonparametric statistical testing of EEG- and MEG-data. J. Neurosci. Methods 164, 177–190. 10.1016/j.jneumeth.2007.03.024

Martin-Ruiz, C., Jagger, C., Kingston, A., Collerton, J., Catt, M., Davies, K., Dunn, M., Hilkens, C., Keavney, B., Pearce, S.H.S., den Elzen, W.P.J., Talbot, D., Wiley, L., Bond, J., Mathers, J.C., Eccles, M.P., Robinson, L., James, O., Kirkwood, T.B.L., von Zglinicki, T., 2011. Assessment of a large panel of candidate biomarkers of ageing in the Newcastle 85+ study. Mech. Ageing Dev. 132, 496–502. 10.1016/j.mad.2011.08.001

Moezzi, B., Pratti, L.M., Hordacre, B., Graetz, L., Berryman, C., Lavrencic, L.M., Ridding, M.C., Keage, H.A.D., McDonnell, M.D., Goldsworthy, M.R., 2019. Characterization of Young and Old Adult Brains: An EEG Functional Connectivity Analysis. Neuroscience 422, 230–239. 10.1016/j.neuroscience.2019.08.038

Musaeus, C.S., Nielsen, M.S., Musaeus, J.S., Høgh, P., 2020. Electroencephalographic Cross-Frequency Coupling as a Sign of Disease Progression in Patients With Mild Cognitive Impairment: A Pilot Study. Front. Neurosci. 14, 544484. 10.3389/fnins.2020.00790

Nagel, B.J., Herting, M.M., Maxwell, E.C., Bruno, R., Fair, D., 2013. Hemispheric lateralization of verbal and spatial working memory during adolescence. Brain Cogn. 82, 58–68. 10.1016/j.bandc.2013.02.007

Niccoli, T., Partridge, L., 2012. Ageing as a risk factor for disease. Curr. Biol. 10.1016/j.cub.2012.07.024

Niemann, H., Sturm, W., Thöne-Otto, A., Willmes, K., 2008. CVLT-California Verbal Learning Test - Deutsche Adaptation. Pearson Assessment.

Nunez, P.L., Srinivasan, R., 2009. Electric Fields of the Brain: The neurophysics of EEG, Electric Fields of the Brain: The neurophysics of EEG. Oxford University Press. 10.1093/acprof:oso/9780195050387.001.0001

Nyberg, L., Salami, A., Andersson, M., Eriksson, J., Kalpouzos, G., Kauppi, K., Lind, J., Pudas, S., Persson, J., Nilsson, L.G., 2010. Longitudinal evidence for diminished frontal cortex function in aging. Proc. Natl. Acad. Sci. U. S. A. 107, 22682–22686. 10.1073/pnas.1012651108

Oostenveld, R., Fries, P., Maris, E., Schoffelen, J.M., 2011. FieldTrip: Open source software for advanced analysis of MEG, EEG, and invasive electrophysiological data. Comput. Intell. Neurosci. 2011. 10.1155/2011/156869

Palmer, J.A., Kreutz-Delgado, K., Rao, B.D., Makeig, S., 2007. Modeling and estimation of dependent subspaces with non-radially symmetric and skewed densities, in: Lecture Notes in Computer Science (Including Subseries Lecture Notes in Artificial Intelligence and Lecture Notes in Bioinformatics). Springer Verlag, pp. 97–104. 10.1007/978-3-540-74494-8_13

Pascual-Marqui, R.D., 2007. Discrete, 3D distributed, linear imaging methods of electric neuronal activity. Part 1: exact, zero error localization. arXiv.

Peltz, C.B., Kim, H.L., Kawas, C.H., 2010. Abnormal EEGs in cognitively and physically healthy oldest old: Findings from the 90+ study. J. Clin. Neurophysiol. 27, 292–295. 10.1097/WNP.0b013e3181eaad7d

Pernet, C.R., Chauveau, N., Gaspar, C., Rousselet, G.A., 2011. LIMO EEG: A toolbox for hierarchical linear modeling of electroencephalographic data. Comput. Intell. Neurosci. 2011. 10.1155/2011/831409

Pion-Tonachini, L., Kreutz-Delgado, K., Makeig, S., 2019. The ICLabel dataset of electroencephalographic (EEG) independent component (IC) features. Data Br. 25, 104101. 10.1016/j.dib.2019.104101

Polich, J., 1997. EEG and ERP assessment of normal aging. Electroencephalogr. Clin. Neurophysiol. - Evoked Potentials 104, 244–256. 10.1016/S0168-5597(97)96139-6

Ray, M.K., Mackay, C.E., Harmer, C.J., Crow, T.J., 2008. Bilateral generic working memory circuit requires left-lateralized addition for verbal processing. Cereb. Cortex 18, 1421–1428. 10.1093/cercor/bhm175

Reid, L.M., MacLullich, A.M.J., 2006. Subjective memory complaints and cognitive impairment in older people. Dement. Geriatr. Cogn. Disord. 10.1159/000096295

Rossiter, H.E., Davis, E.M., Clark, E. V., Boudrias, M.H., Ward, N.S., 2014. Beta oscillations reflect changes in motor cortex inhibition in healthy ageing. Neuroimage 91, 360–365. 10.1016/j.neuroimage.2014.01.012

Sahoo, B., Pathak, A., Deco, G., Banerjee, A., Roy, D., 2020. Lifespan associated global patterns of coherent neural communication. Neuroimage 216, 116824. 10.1016/j.neuroimage.2020.116824

Scally, B., Burke, M.R., Bunce, D., Delvenne, J.F., 2018. Resting-state EEG power and connectivity are associated with alpha peak frequency slowing in healthy aging. Neurobiol. Aging 71, 149–155. 10.1016/j.neurobiolaging.2018.07.004

Schmidt, B.T., Ghuman, A.S., Huppert, T.J., 2014. Whole brain functional connectivity using phase locking measures of resting state magnetoencephalography. Front. Neurosci. 8, 141. 10.3389/fnins.2014.00141

Schnitzler, A., Gross, J., 2005. Normal and pathological oscillatory communication in the brain. Nat. Rev. Neurosci. 10.1038/nrn1650

Schoonhoven, D.N., Briels, C.T., Hillebrand, A., Scheltens, P., Stam, C.J., Gouw, A.A., 2022. Sensitive and reproducible MEG resting-state metrics of functional connectivity in Alzheimer’s disease. Alzheimer’s Res. Ther. 14, 1–19. 10.1186/s13195-022-00970-4

Siebenhühner, F., Wang, S.H., Arnulfo, G., Lampinen, A., Nobili, L., Palva, J.M., Palva, S., 2020. Genuine cross-frequency coupling networks in human resting-state electrophysiological recordings. PLoS Biol. 18, e3000685. 10.1371/journal.pbio.3000685

Simpraga, S., Alvarez-Jimenez, R., Mansvelder, H.D., Van Gerven, J.M.A., Groeneveld, G.J., Poil, S.S., Linkenkaer-Hansen, K., 2017. EEG machine learning for accurate detection of cholinergic intervention and Alzheimer’s disease. Sci. Rep. 7, 1–11. 10.1038/s41598-017-06165-4

Taylor, J.R., Williams, N., Cusack, R., Auer, T., Shafto, M.A., Dixon, M., Tyler, L.K., Cam-CAN, Henson, R.N., 2017. The Cambridge Centre for Ageing and Neuroscience (Cam-CAN) data repository: Structural and functional MRI, MEG, and cognitive data from a cross-sectional adult lifespan sample. Neuroimage 144, 262–269. 10.1016/j.neuroimage.2015.09.018

Tooley, U.A., Park, A.T., Leonard, J.A., Boroshok, A.L., McDermott, C.L., Tisdall, M.D., Bassett, D.S., Mackey, A.P., 2022. The Age of Reason: Functional Brain Network Development during Childhood. J. Neurosci. 42, 8237–8251. 10.1523/JNEUROSCI.0511-22.2022

Trammell, J.P., MacRae, P.G., Davis, G., Bergstedt, D., Anderson, A.E., 2017. The relationship of cognitive performance and the Theta-Alpha power ratio is age-dependent: An EEG study of short term memory and reasoning during task and resting-state in healthy young and old adults. Front. Aging Neurosci. 9, 364. 10.3389/FNAGI.2017.00364/BIBTEX

Tröndle, M., Popov, T., Pedroni, A., Pfeiffer, C., Barańczuk-Turska, Z., Langer, N., 2023. Decomposing age effects in EEG alpha power. Cortex 161, 116–144. 10.1016/j.cortex.2023.02.002

Van Der Hiele, K., Bollen, E.L.E.M., Vein, A.A., Reijntjes, R.H.A.M., Westendorp, R.G.J., Van Buchem, M.A., Middelkoop, H.A.M., Van Dijk, J.G., 2008. EEG markers of future cognitive performance in the elderly. J. Clin. Neurophysiol. 25, 83–89. 10.1097/WNP.0b013e31816a5b25

Veldhuizen, R.J., Jonkman, E.J., Poortvliet, D.C.J., 1993. Sex differences in age regression parameters of healthy adults-normative data and practical implications. Electroencephalogr. Clin. Neurophysiol. 86, 377–384. 10.1016/0013-4694(93)90133-G

Vinck, M., Oostenveld, R., Van Wingerden, M., Battaglia, F., Pennartz, C.M.A., 2011. An improved index of phase-synchronization for electrophysiological data in the presence of volume-conduction, noise and sample-size bias. Neuroimage 55, 1548–1565. 10.1016/j.neuroimage.2011.01.055

Vlahou, E.L., Thurm, F., Kolassa, I.T., Schlee, W., 2014. Resting-state slow wave power, healthy aging and cognitive performance. Sci. Rep. 4, 1–6. 10.1038/srep05101

Vorwerk, J., Oostenveld, R., Piastra, M.C., Magyari, L., Wolters, C.H., 2018. The FieldTrip-SimBio pipeline for EEG forward solutions. Biomed. Eng. Online 17, 37. 10.1186/s12938-018-0463-y

Voytek, B., Kramer, M.A., Case, J., Lepage, K.Q., Tempesta, Z.R., Knight, R.T., Gazzaley, A., 2015. Age-related changes in 1/f neural electrophysiological noise. J. Neurosci. 35, 13257–13265. 10.1523/JNEUROSCI.2332-14.2015

Vysata, O., Kukal, J., Prochazka, A., Pazdera, L., Valis, M., 2012. Age-related changes in the energy and spectral composition of EEG. Neurophysiology 44, 63–67. 10.1007/s11062-012-9268-y

Wan, L., Huang, H., Schwab, N., Tanner, J., Rajan, A., Lam, N.B., Zaborszky, L., Li, C. shan R., Price, C.C., Ding, M., 2019. From eyes-closed to eyes-open: Role of cholinergic projections in EC-to-EO alpha reactivity revealed by combining EEG and MRI. Hum. Brain Mapp. 40, 566–577. 10.1002/hbm.24395

Zhang, Y., Wu, W., Toll, R.T., Naparstek, S., Maron-Katz, A., Watts, M., Gordon, J., Jeong, J., Astolfi, L., Shpigel, E., Longwell, P., Sarhadi, K., El-Said, D., Li, Y., Cooper, C., Chin-Fatt, C., Arns, M., Goodkind, M.S., Trivedi, M.H., Marmar, C.R., Etkin, A., 2021. Identification of psychiatric disorder subtypes from functional connectivity patterns in resting-state electroencephalography. Nat. Biomed. Eng. 5, 309–323. 10.1038/s41551-020-00614-8

Zhong, X., Chen, J.J., Chen, J., 2020. Variations in the frequency and amplitude of resting-state EEG and fMRI signals in normal adults: The effects of age and sex. bioRxiv 2020.10.02.323840. 10.1101/2020.10.02.323840

Ziemmermann, P., Fimm, B., 2021. A test battery for attentional performance, in: Applied Neuropsychology of Attention. Psychology Press, pp. 124–165. 10.4324/9780203307014-12

Ziemmermann, P., Fimm, B., n.d. Psytest - TAP 2.3.1 [WWW Document]. URL https://www.psytest.de/index.php?page=TAP-2-2&hl=de_DE (accessed 6.24.21).

Zou, Q., Ross, T.J., Gu, H., Geng, X., Zuo, X.N., Hong, L.E., Gao, J.H., Stein, E.A., Zang, Y.F., Yang, Y., 2013. Intrinsic resting-state activity predicts working memory brain activation and behavioral performance. Hum. Brain Mapp. 34, 3204–3215. 10.1002/hbm.22136

