## Supplementary figures and images for "Functionally annotated electrophysiological neuromarkers of healthy ageing and memory function"

### Supplemental figure 1

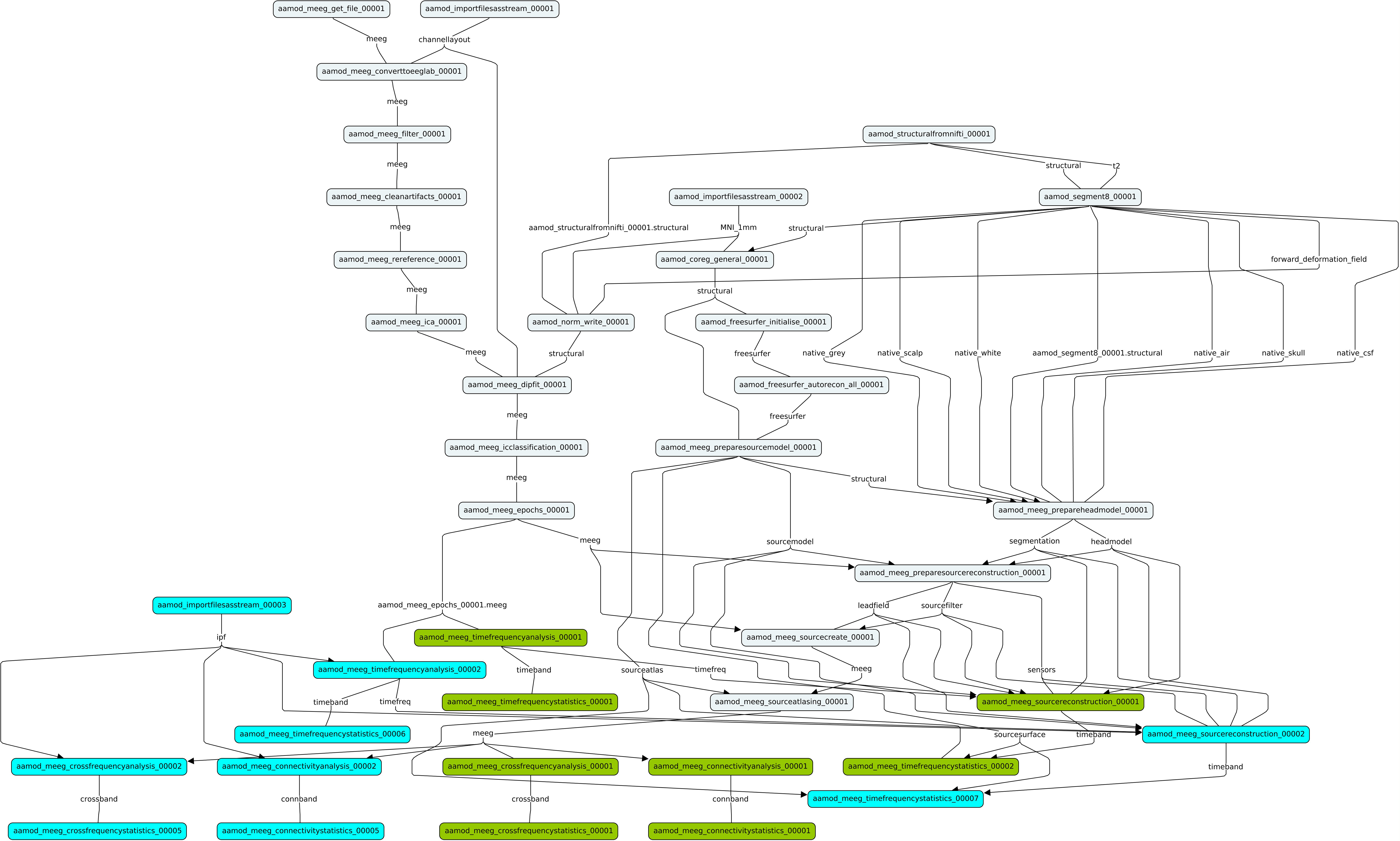

### Supplemental figure 2

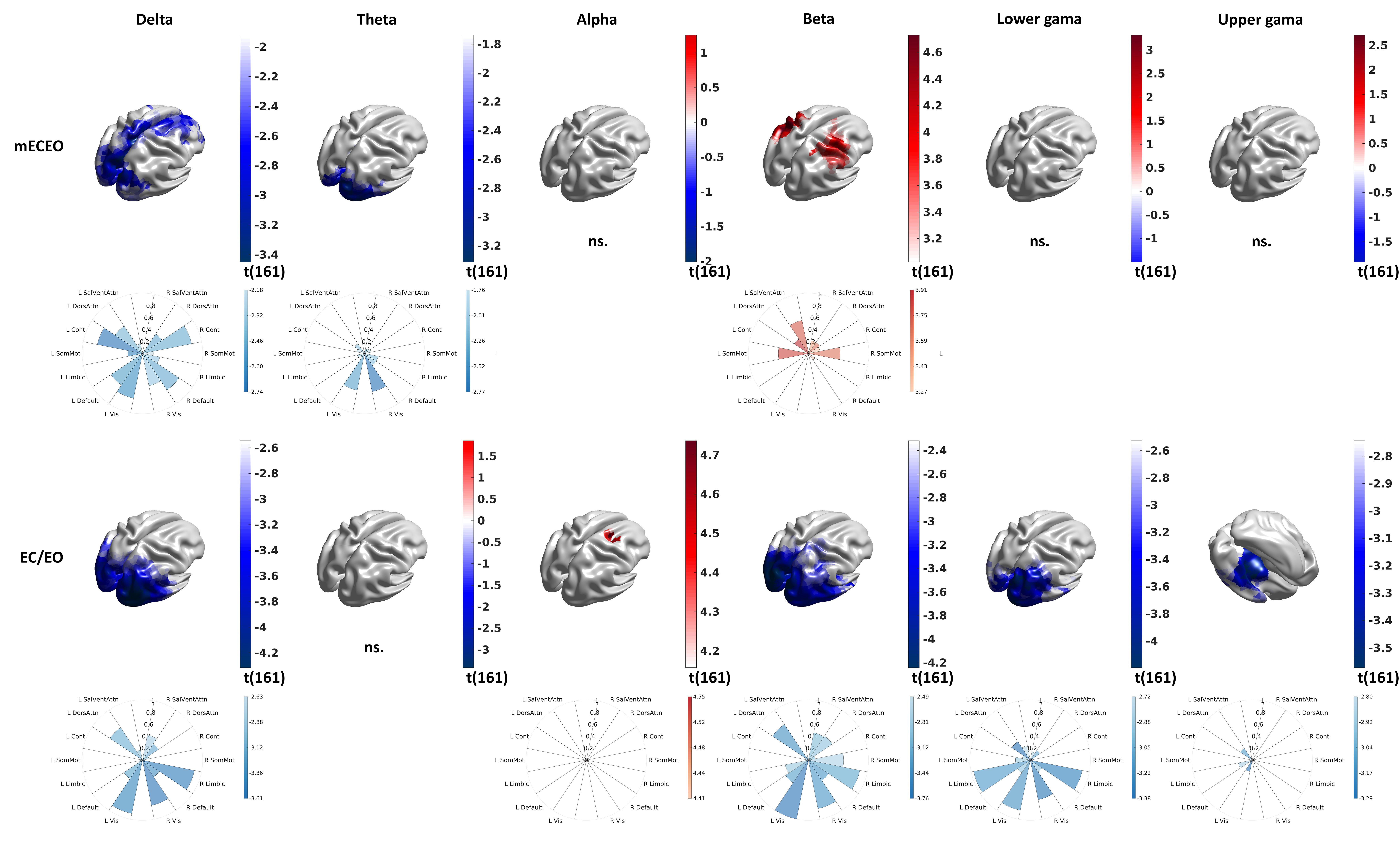

### Supplemental figure 3

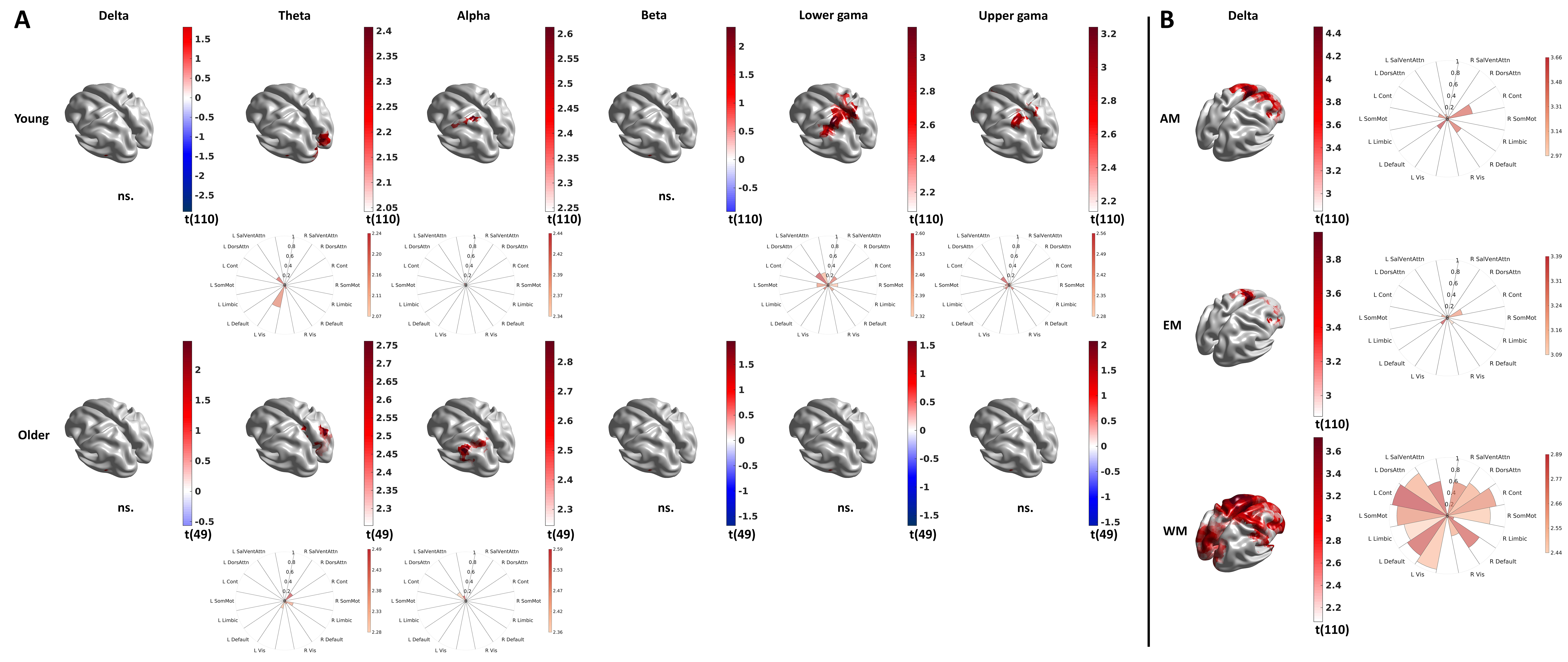
